# Priority effects of heritable seed-borne bacteria drive early assembly of the wheat rhizosphere microbiome

**DOI:** 10.1101/2024.10.21.619384

**Authors:** Daniel Garrido-Sanz, Christoph Keel

## Abstract

Microbial communities play a critical role in supporting plant health and productivity, making the ability to obtain reproducible plant-associated microbiomes an essential asset for experimentally testing hypotheses related to microbiome manipulation and fundamental principles governing community dynamics. We used a sequential propagation strategy to generate a complex and reproducible wheat rhizosphere microbiome (RhizCom) that was shaped by host selection and periodic habitat resetting. Heritable seed-borne rhizosphere bacteria (SbRB) emerged as the dominant microbiome source after coalescing with the soil community, driven by priority effects and efficient niche exploitation during early habitat development. Functional analyses revealed that niche partitioning through the ability of SbRB to degrade specific saccharides and niche facilitation contributed to the assembly of the RhizCom. Our results advance our understanding of the principles governing microbial community dynamics in early plant development and provide strategies for future microbiome manipulation aimed at improving crop productivity and health.

## Introduction

Plant-associated microbiomes carry a wealth of functions that are intrinsically related to the growth and health of the host^1–4^ and collectively contribute to the functioning of the plant holobiont^5^. In agroecosystems, the rhizosphere microbiome (i.e., rhizobiome) plays a critical role in crop productivity^6,7^, and approaches to manipulate it hold great promise for meeting increasing agricultural demands without resorting to harmful chemical fertilizers and pesticides.

Targeted microbiome manipulation focuses on the addition, removal or modulation of specific functions within the rhizobiome, achieved through the introduction of plant-beneficial inoculants or the elimination of harmful pathogens^8^. This includes bacteria capable of solubilizing plant growth-limiting nutrients^1^, modulating plant stress responses to tolerate moderate drought^2^, antagonizing phytopathogens^3^ or priming plant responses to insect pests^9^, or using highly specific bacteriophages to reduce a target population^10^. However, the success of plant microbiome engineering is often constrained by a limited understanding of how complex communities respond to such interventions^11,12^. This is not surprising, as soil and rhizosphere microbiomes are among the most diverse environments on Earth^13,14^, making them particularly challenging for mechanistic studies. Bottom-up approaches using synthetic communities (SynComs) offer highly tractable and reproducible systems in which complex interaction mechanisms can be elucidated^15–17^, yet they often represent an oversimplification of natural environments. In contrast, top-down strategies consist of the generation of natural communities (NatComs) by culturing microbiomes in controlled set-ups that mimic natural environments^18–20^, allowing the selection of complex microbiomes that closely resemble the structure and composition of their original environment. Despite their potential, the widespread adoption of NatComs remains limited. This is largely due to their inherent biological complexity, the lack of reproducible natural microbial communities, and the limitations of current technological and analytical methods.

The selection exerted by root exudates is arguably one of the main factors determining the rhizobiome assembly. Root exudates are composed of carbon-rich molecules, predominantly organic acids, amino acids and saccharides, which support net bacterial growth in high numbers^21,22^. In addition, root exudates contain signaling molecules that control the proliferation of specific bacterial taxa^23–25^. Owing to the diffusion of root exudates into the adjacent soil, soil-dwelling bacteria are inherently considered the major source of microbes colonizing the rhizosphere. However, recent studies have emphasized that seed-transmitted bacteria (STB) are also an important source of the rhizobiome^26–28^.

These heritable bacteria are found across many crops and agriculturally relevant plants^29^. STB reside in the internal tissues of seeds, which they reach by different transmission routes^30^. Once the seed germinates, STB may leave the seed tissues and become part of the new plant rhizobiome, i.e. seed-borne rhizosphere bacteria (SbRB). These bacteria contribute to beneficial functions for the plant^31,32^, especially during early stages of plant development^33^, independent of their availability in the surrounding soil. However, how both SbRB and soil microbiomes coalesce into the plant rhizobiome and whether specific metabolic signatures exist within the heritable plant microbial diversity remains poorly investigated.

In this study, we used a sequential wheat rhizobiome propagation approach to obtain a stable and reproducible species-rich rhizosphere natural community (RhizCom). This allowed us to investigate the successional dynamics and coalescence of the initial soil microbiome and SbRB, leading to the assembly of a reproducible wheat rhizobiome constrained by plant root selection. Functional analyses revealed the enrichment of plant-interaction traits in the root-associated communities, as opposed to the broader metabolic diversity observed in the soil community. The specific metabolism of saccharides in SbRB suggests their critical role in facilitating the assembly of the rhizosphere microbiome. This work provides an unprecedented view of early rhizosphere microbiome assembly in the world’s third most productive crop, facilitating the obtention of reproducible and complex bacterial communities aimed for future mechanistic studies.

## Results

### Sequential succession of the wheat rhizobiome drives phylum-specific selection

To obtain a tractable wheat rhizobiome, we performed six successive cycles of microbiome propagation in the rhizosphere of wheat (*Triticum aestivum* cv. Arina) using a soil microbiome cell suspension as the initial inoculum. After seven days of plant development, the rhizosphere was collected, and a rhizosphere cell suspension was obtained and used to reinoculate the next cycle (Fig. 1a, Extended Data Fig. 1). We tracked bacterial community dynamics based on analysis of the small ribosomal subunit (16S) rRNA gene. A strong shift from the initial soil microbiome composition was observed, caused by the selection of bacteria mainly belonging to the phyla *Proteobacteria* and *Bacteroidota*, and to a lesser extent some members of *Firmicutes* and *Verrucomicrobiota* (Fig. 1b, Supplementary Fig. 1). SbRB and soil samples were the most dissimilar compared to the microbiome composition observed along the succession cycles, which showed a converging trajectory (Fig. 1cd). Taxa selected in the initial cycles were maintained throughout the remaining steps (Fig. 1e, Supplementary Fig. 2), consistent with a stabilization of phylogenetic diversity (Spearman correlation; R = −0.15, *P* value = 0.49) and colony-forming unit (c.f.u.) counts (Fig. 1fg, Supplementary Table 1). However, a progressive decrease in Shannon diversity (Spearman correlation; *R* = −0.85, *P* value = 1.2e^-7^, Fig. 1h) and other alpha diversity indices was observed (Supplementary Fig. 3). These data demonstrate the establishment of a stable and diverse wheat rhizosphere community (RhizCom) under the microcosm conditions used, consisting of 464 ASVs belonging to 126 distinct genera (Supplementary Table 2). The reproducibility of the RhizCom was validated by re-growing the cryopreserved RhizCom at −80 °C in the wheat rhizosphere. The recovered RhizCom had a similar taxonomic composition, number of ASVs, and Shannon diversity as the community of the last propagation cycle (Fig. 1, Supplementary Fig. 3, Supplementary Table 3).

**Fig. 1.**
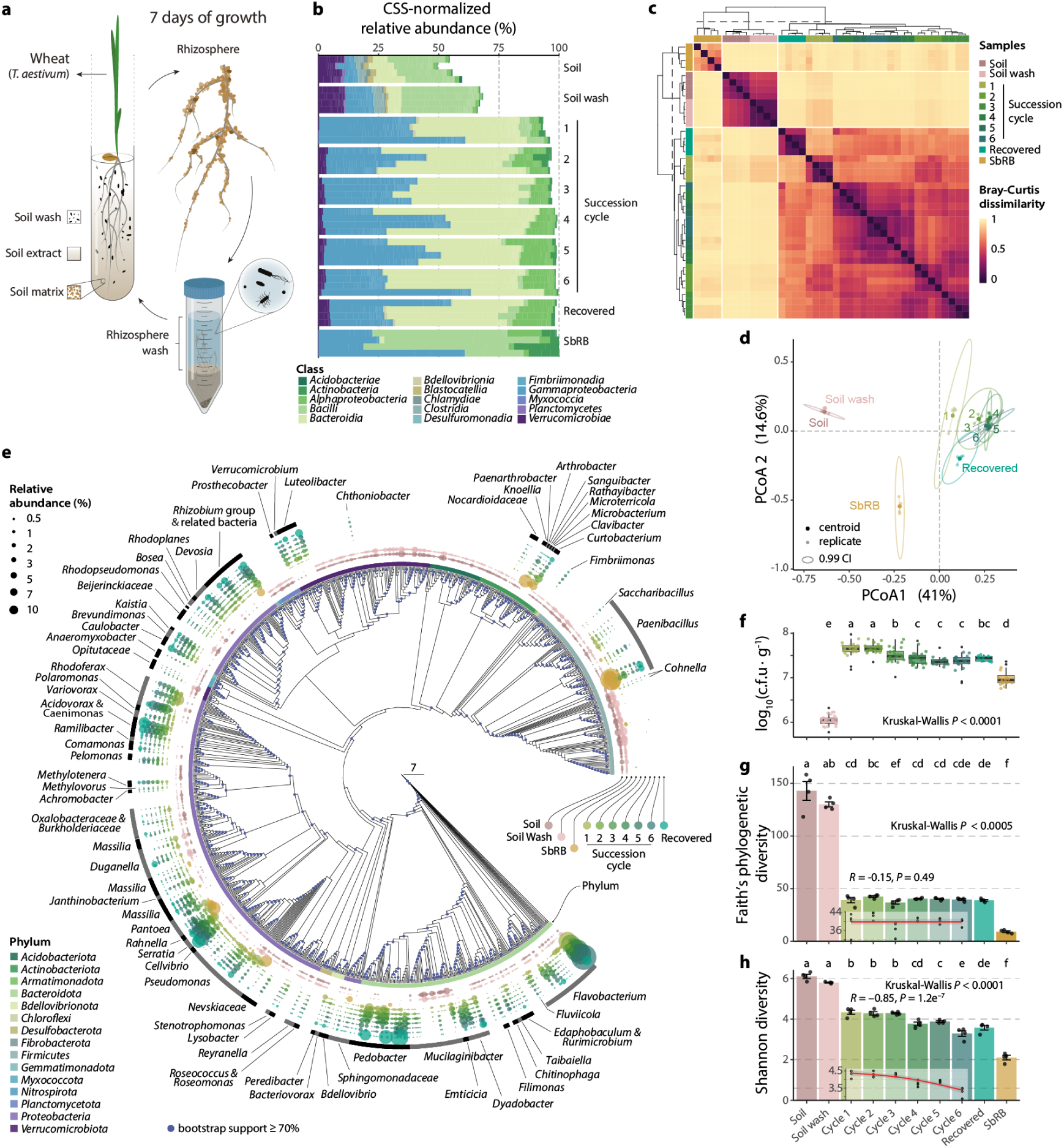
Sequential succession of a wheat rhizosphere microbiome. **a**, Schematic representation of the microcosm used and experimental design. See the Extended Data Fig. 1 for details. **b**, CSS-normalized relative abundance of the top 500 ASVs at the class level. **c**, Clustering of Bray-Curtis dissimilarities across sample replicates (*n* = 4 per sample) showed a clear distinction of SbRB, soil, soil wash and succession cycles. **d**, Principal coordinate analysis (PCoA) based on Bray-Curtis dissimilarities across samples. The ellipses represent a 99% confidence interval. **e**, Selection of ASVs during the succession of the wheat rhizobiome. The phylogenetic tree was built using ASVs with a total mean (*n* = 4) relative abundance ≥ 0.005%. Only the names of relevant genera selected in the rhizosphere microbiome succession are indicated. From inwards to outwards, colored dots represent ASVs, with sizes corresponding to their mean relative abundance in samples. For details see Supplementary Fig. 2. **f**, c.f.u. enumeration of bacteria on R-2A agar per gram of soil (soil wash, *n* = 28) or rhizosphere (*n* >15, recovered *n* = 8). **g,h**, Measures of alpha diversity across samples and Spearman correlation with the succession cycles. The mean values are represented by bars (± standard deviation). Each black dot represents a replicate (*n* = 4 per sample). Different letters indicate statistically significant groups (*P* value ≤ 0.01) determined using Kruskal-Wallis test, followed by Fisher’s post hoc test and fdr *P* value adjustment. The inner plots show the Spearman correlation between alpha diversity measurements and succession cycles (red line; mean values, dots; replicates (*n* = 4)). Correlation coefficient (*R*) and *P* value (*P*) are indicated.

### Heritable SbRB constitute the major rhizobiome source

We examined the contribution of the soil wash inoculum and SbRB to the RhizCom assemblage. Only a small number of RhizCom ASVs could be traced to either the SbRB or the soil wash source communities (11.4%, Fig. 2a, Supplementary Table 3). Specifically, we detected 33 ASVs originating from the soil inoculum and 25 from the SbRB in the RhizCom microbiome, which together accounted for 51% of the total RhizCom relative abundance. Notably, one *Flavobacterium* ASV and one *Serratia* ASV, both detected in the soil inoculum and the SbRB community, accounted for 43.3% of the RhizCom (Supplementary Table 3). Soil wash and SbRB ASVs contributing to the RhizCom collectively represented 4% and 84.7% of the relative abundance of their respective microbiomes. Most ASVs not traced to the source communities emerged throughout the succession cycles (70.9% of RhizCom ASVs), indicating that individual heritable SbRB heterogeneity was retained within the RhizCom (Supplementary Table 3). This was further evidenced by a decrease in ASV emergence over the cycles, mostly affecting low abundant taxa (Fig. 2b). RhizCom ASVs not detected in any condition (82 ASVs, Fig. 2a) had a low representation in the RhizCom (85% of them below 0.01% of relative abundance, Supplementary Table 3) and could be attributed to specific SbRB heterogeneity within its cycle.

**Fig. 2.**
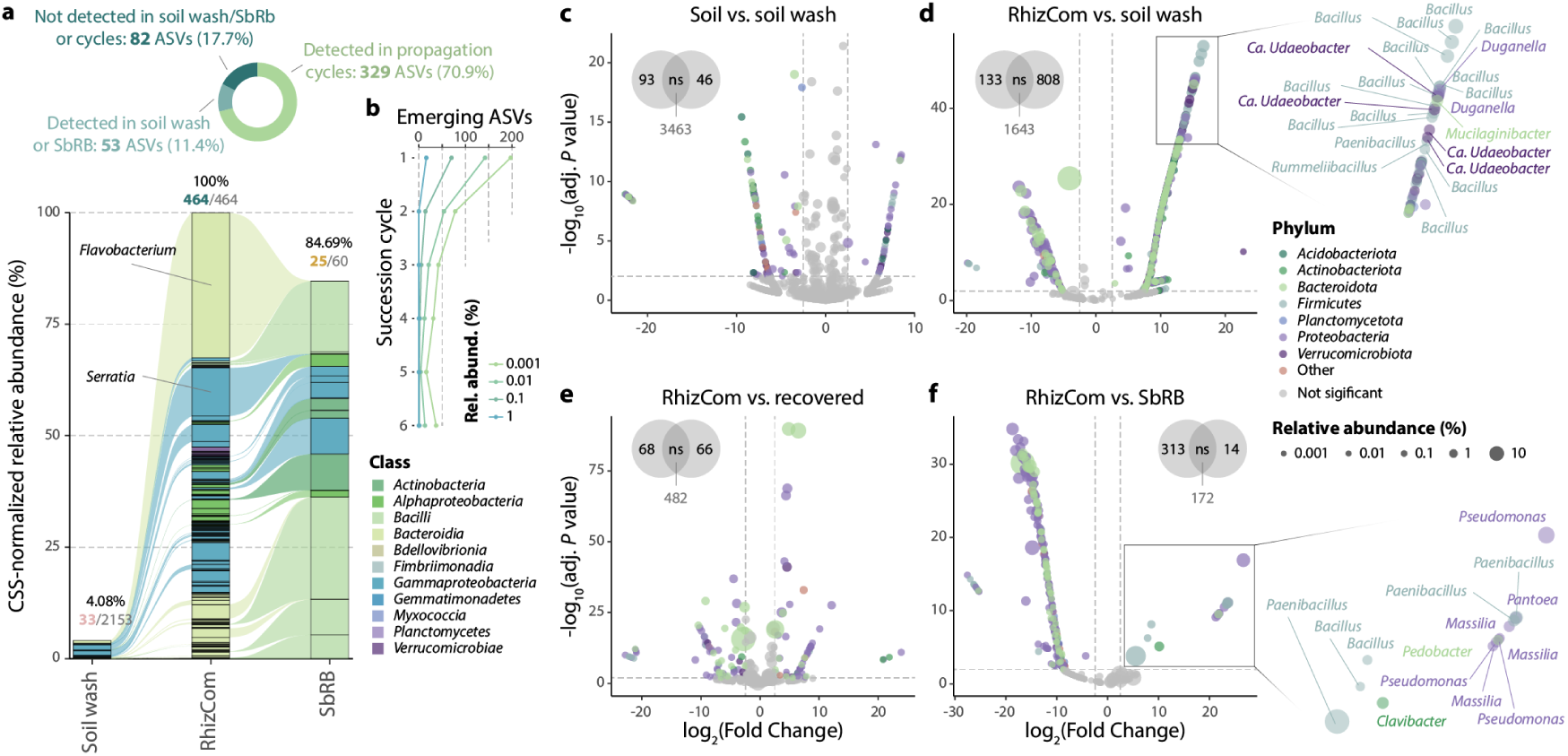
Contribution of SbRB to the RhizCom. **a**, RhizCom ASVs traced to the initial soil wash inoculum or the seed-borne rhizosphere bacteria (SbRB). Alluvial diagram shows the connection of ASVs throughout the three samples. Only ASVs present in the RhizCom (6^th^ succession cycle) are considered. Replicates (*n* = 4) were pooled by mean values. Percentages show the combined CSS-normalized relative abundance of the ASVs that contribute to the RhizCom, followed by the number of ASVs / total ASVs in the sample in bold / grey according to the sample, respectively. Color according to ASVs taxonomy at the class level. The doughnut chart on top shows the number of the RhizCom ASVs detected in the soil wash or the SbRB. **b**, Emergence of ASVs through the succession cycles. Lines represent the number of new ASVs detected at each cycle at different thresholds of relative abundance. **c-f**, Differential ASV abundance calculated with DESeq2 using the Wald test for significance and a local estimate of dispersion. ASVs with a |log2(fold change)| ≥ 2.5 and an adjusted (adj.) *P* value < 0.01 were considered as significant. Results are represented as volcano plots of key comparisons. Dots represent ASVs colored by phylum and sized according to their relative abundance. The number of differentially abundant ASVs per condition is shown within Venn diagrams (ns; not significant). To the right of panels **c** and **e**, zoom with labels on the most depleted ASVs in the RhizCom compared to soil wash or SbRB samples. See Supplementary Fig. 4 for details.

The comparison between the initial soil wash inoculum and the RhizCom revealed the most striking changes, with 808 ASVs enriched in the soil wash vs. 133 in the RhizCom (Fig. 2c-f), followed by the comparison SbRB vs. RhizCom (14 vs. 313 ASVs, respectively). This indicates substantial differences in bacterial community composition, suggesting a strong host-mediated selective pressure. The enrichment of ASVs in the RhizCom mainly affected taxa belonging to *Proteobacteria* and *Bacteroidota*, as also observed during the propagation cycles (Fig. 1f), while ASVs belonging to *Firmicutes*, especially *Bacillus*, remained more enriched in the soil inoculum (Supplementary Fig. 4). An exception were certain SbRB ASVs (*Pseudomonas*, *Massilia* and *Pantoea*) whose relative abundances were reduced compared to the RhizCom (Fig. 2f). This could be attributed to the inability of certain SbRB to maintain a similar abundance in the RhizCom as a result of the coalescence of the two microbiomes. The changes observed between the RhizCom and the recovered community (Fig. 2e) are largely attributable to shifts in relative abundances rather than loss of taxa (Fig. 1b, Supplementary Table 3), further emphasizing that the overall RhizCom taxonomic composition remains stable after recovery (Fig. 1g).

### The wheat rhizobiome and its source communities differ in global functional patterns

The RhizCom, SbRB and initial soil communities were functionally characterized by metagenomic shotgun sequencing. Over 1 billion reads were co-assembled into 1.5 million contigs (> 1,000 bp), totaling 5.2 billion bp (Extended Data Fig. 2, Supplementary Table 4). Sample coverage based on sequence diversity showed nearly complete coverage of the RhizCom and SbRB metagenomes (average coverage > 97%), whereas soil coverage was > 52%, consistent with higher soil sequence diversity (Extended Data Fig. 2b). The samples showed distinct taxonomic profiles dominated by Bacteria (Supplementary Fig. 5), as determined by the number of reads mapped to coding DNA sequences. Among them, *Proteobacteria* were the most prevalent, representing more than 50% in RhizCom and SbRB samples, but about 20% in the soil. Overall, the taxonomic composition of the metagenomic samples was consistent with that obtained by 16S rRNA amplicon sequencing (Supplementary Fig. 6). The communities differed in their general functional content (ANOSIM, *P* value = 0.001, Extended Data Fig. 3), and showed a predominance of genes involved in general cellular functions, including genetic information processing and carbohydrate and amino acid metabolism.

Analysis of the differential abundance of KEGG annotations among the communities and their classification into dominance categories revealed a similar number of enriched functions in both the RhizCom and soil communities (1,989 and 2,012, respectively, Fig. 3a). Enriched functions in the RhizCom were involved in cellular processes, including transport and catabolism, and information processing (Fig. 3b). In contrast, the soil was enriched in functions related to signaling molecules and interaction, and metabolism, including biosynthesis of secondary metabolites, energy metabolism, metabolism of terpenoids and polyketides and biodegradation and metabolism of xenobiotics. SbRB were enriched in membrane transport. Notably, functions with balanced distributions across samples had higher mean abundances than other dominance categories (Fig. 3c, Supplementary Table 5). The enrichment of a category in a given community was not caused by an overall increase in abundance within that community, but rather resulted from lower abundance in the other two communities (Fig. 3c).

**Fig. 3.**
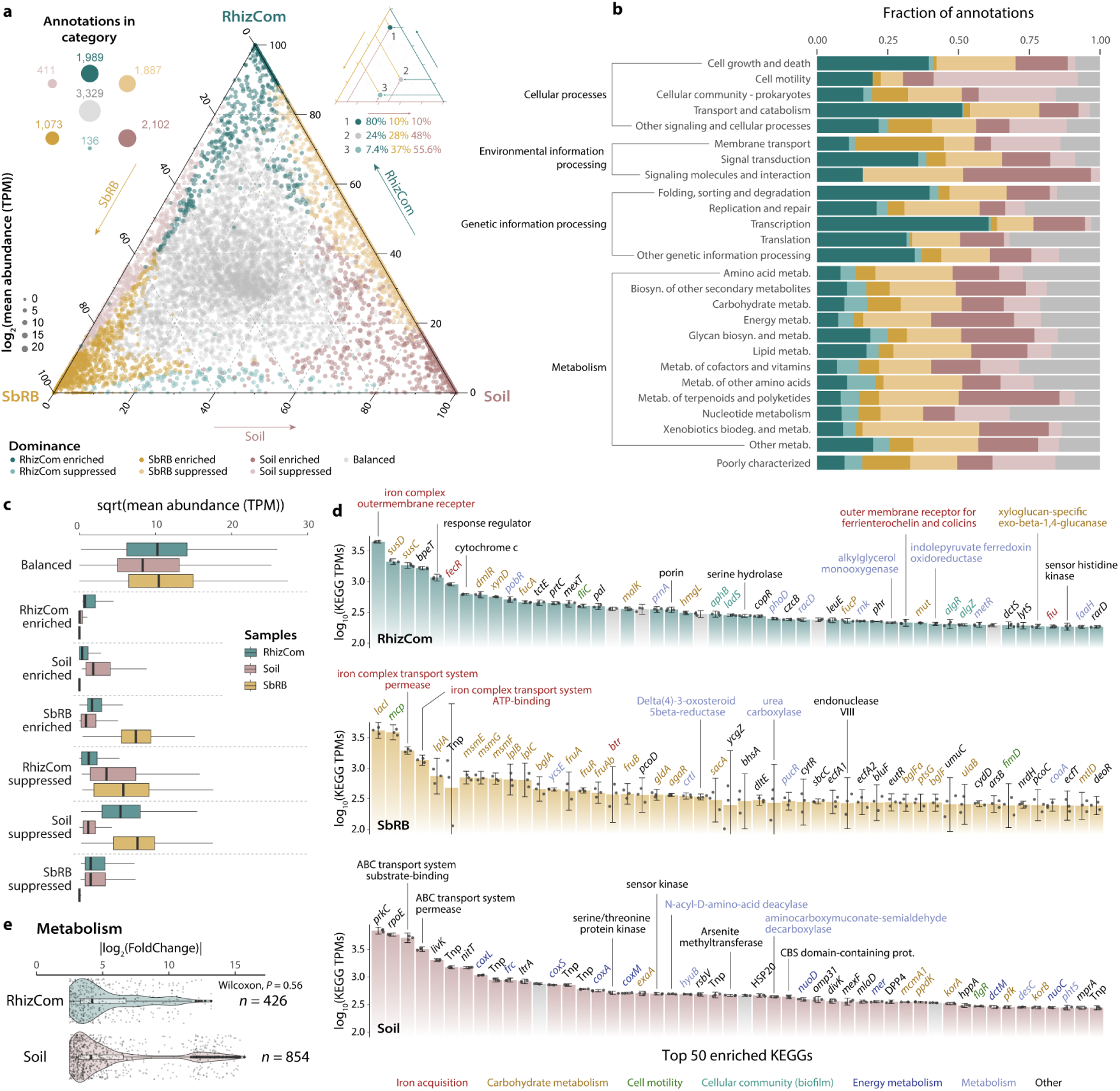
Functional contribution of communities. **a**, Ternary plot showing the contribution of each sample to KEGG functional annotation abundances. Each coordinate (represented by dots) consists of the relative contribution of each function in ternary space. Functions are sized according to log2 of the mean TPM across samples. Only annotations with mean TPM > 0.5 across samples were considered. Color according to dominance categories defined by the result of pairwise differential abundance analyses. Top left: circles and numbers represent the number of functions per dominance category, with sizes at scale. Top right: example of interpretation of function contribution in the ternary space. **b**, Fraction of annotations assigned to the defined dominance categories in panel **a** per functional KEGG BRITE hierarchy. **c**, Square root (sqrt) of mean KEGG TPMs per sample (*n* = 3 replicates per sample) and dominance category. Statistical differences are shown in the Supplementary Table 5. **d**, Abundance (TPMs) of the top 50 functions in samples’ enriched categories. Bars represent mean values (± standard error). Black dots represent individual replicate values (*n* = 3). Gene names or annotations are indicated. Grey bars indicate KEGGs with unknown functions. Full information across all dominance categories is provided in Supplementary Table 6. **e**, Differentially abundant (|log2(fold change)| ≥ 2.5 and an adjusted *P* value < 0.01) KEGG annotations involved in metabolism between RhizCom and soil. Statistical differences were calculated using the Wilcoxon test. The number of differentially abundant annotations is indicated, and they show a higher value in the soil, although the change in abundance (i.e., fold change) remains similar (*P* value = 0.56).

### The wheat rhizobiome is enriched in host-interaction traits

When considering the most enriched functions, iron acquisition was a dominant feature in both the RhizCom and SbRB communities (Fig. 3d). Enriched functions in the RhizCom also included genes for starch and sucrose metabolism, and plant colonization and beneficial activities such as the biosynthesis of flagella, exopolysaccharides, alkaline phosphatase, hydrogen cyanide, and 1-aminocyclopropane-1-carboxylate (ACC) deaminase (Fig. 3d, Supplementary Table 6). The enriched functions in SbRB included genes involved in the transport and metabolism of different saccharides, including cellulose/hemicellulose intermediates. The soil community was enriched in broader substate metabolism and energy production, including the metabolism of sulfur and methane, and the biodegradation of aromatic compounds, including toluene, xylene, phthalate, and dioxins (Fig. 3d, Supplementary Table 6). Furthermore, since SbRB was predominantly integrated into the RhizCom (Fig. 2a), we examined the differential abundance of KEGG annotations between the RhizCom and the soil (Supplementary Table 7). The results emphasized the enrichment of genes involved in biofilm formation, motility and carbohydrate metabolism in the RhizCom, whereas the soil was characterized by genes involved in the biodegradation and metabolism of a wide range of compounds (Fig. 3e, Supplementary Fig. 7).

Enriched functions followed a distinct taxonomic distribution within the communities (Extended Data Fig. 4), reflecting the dominant members of each community. In the SbRB, most enriched functions resided in members of *Actinomycetia*, *Bacilli* and *Gammaproteobacteria*, particularly in genera such as *Pantoea*, *Paenibacillus*, *Curtobacterium, Sanguibacter, Clavibacter*, and members of the *Rhizobium* group. Conversely, enriched functions in the RhizCom were mainly associated with *Proteobacteria* and *Bacteroidota*, especially *Acidovorax, Flavobacterium*, *Pedobacter*, *Caulobacter*, and *Pseudomonas*. In the soil community, enriched functions were more broadly distributed across mostly unclassified bacteria at various taxonomic ranks.

### Metabolic specialization of heritable SbRB drives saccharide metabolism in the wheat rhizobiome

Binning of contigs and dereplication of bins resulted in a total of 821 metagenome-assembled genomes (MAGs), of which 110 were of high to medium quality (completeness >50%, contamination <10%, Fig. 4a). These MAGs represented 22.2% of the RhizCom, 42.8% of the SbRB, and 8.4% of the soil communities (Fig. 4b, Supplementary Table 8). The assignment of MAGs to their primary community based on TPMs resulted in 60 MAGs being assigned to the RhizCom, 16 to the SbRB, and 33 to the soil communities (Fig. 4c, Supplementary Table 8). The taxonomic classification of the MAGs corresponds to the most abundant taxa on each community. For example, MAGs assigned to SbRB belong to *Pantoea*, *Paenibacillus*, *Rhizobium* or *Pseudomonas* (Fig. 4c, Fig. 1e).

**Fig. 4.**
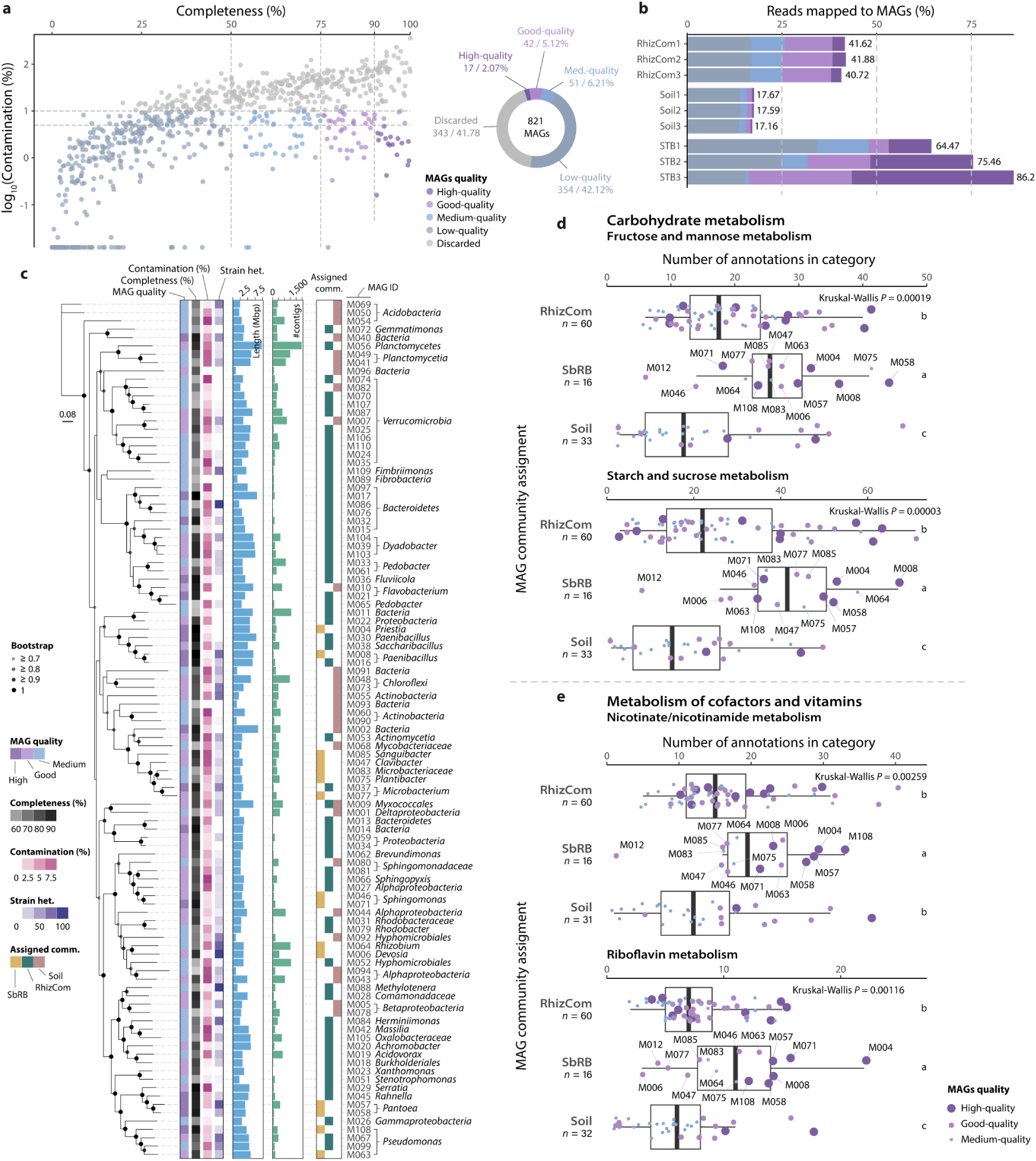
Characteristics of metagenome-assembled genomes (MAGs). **a**, MAG quality based on percentage of completeness and contamination. High quality MAGs: > 90% completeness, < 5% contamination; good-quality MAGs: > 75% completeness, < 10% contamination; medium quality MAGs: < 50% completeness, < 10% contamination; low quality MAGs: ≤ 50% contamination, < 10% contamination; discarded MAGs: ≥ 10% contamination. Dashed grey lines indicate the MAG quality thresholds. The number and percentage of MAGs per quality category are shown in the doughnut plot. **b**, Percentage of the total number of reads mapped to MAGs. Unmapped and unclassified reads were not considered for calculation of percentages. **c**, Taxonomic classification of high to medium quality MAGs and characteristics. Taxonomic classification was based on the last taxonomic rank as obtained by MAG contig consensus. Phylogenetic tree based on maximum-likelihood (ML) of concatenated conserved amino acid markers, expanding 38,640 aligned positions, and using 100 rapid bootstrap inferences followed by a thorough ML search for the best scoring tree. Seven MAGs lacking sufficient marker gene sequences for phylogenetic inference were discarded. **d**,**e**, Metabolic features enriched in SbRB MAGs. Number of annotations in fructose and mannose and starch and sucrose (**d**) or nicotinate/nicotinamide (**e**) metabolic categories.

Functional differences between the MAGs assigned to the three communities were explored based on KEGG annotations. The results show a higher proportion of annotations for starch and sucrose metabolism (Kruskal-Wallis: *P* value = 0.00003) and fructose and mannose metabolism (*P* value = 0.00019) in SbRB MAGs (Fig. 4d), as well as a higher number of annotations involved in nicotinate and nicotinamide metabolism (*P* value = 0.00259) and riboflavin metabolism (*P* value = 0.00116), suggesting a specialized SbRB metabolism. SbRB with the highest number of annotations in these categories belong to *Pantoea* (MAGs M057 and M058), *Paenibacillus* (M008), *Pseudomonas* (M063 and M108) and *Priestia* (M004), consistent with their prevalence (Fig. 1e, Extended Fig. 5, Supplementary Tables S3 and S8).

Further evaluation of specific reactions differentially present in the MAGs of each community revealed that SbRB are capable of assimilating specific extracellular disaccharides: sucrose, maltose, cellobiose and trehalose via their conversion to the more common intermediate monosaccharides fructose and glucose (Fig. 5). While these upper pathways were absent in members of the soil community, enzymes for lower metabolic pathways (e.g. conversion of D-glucose to D-fructose-6P, or D-fructose to D-fructose-6P, Fig. 5b) were similarly encoded by MAGs from the three communities (Fig. 5c). In addition, MAGs assigned to SbRB and RhizCom showed a similar prevalence of upper disaccharide assimilation pathways. Most of these reactions were assigned to the most abundant members of SbRB: *Pantoea*, *Paenibacillus* and *Priestia*. Additionally, complete pathways for the biosynthesis of nicotinate and riboflavin vitamins (B_3_ and B_2_, respectively) were also more prevalent in SbRB MAGs than in soil (6 vs. 4 for nicotinate and 8 vs. 3 for riboflavin, respectively), with both pathways present in *Pantoea*, *Paenibacillus*, *Priestia* and *Pseudomonas* SbRB MAGs (Fig. 5a). These results suggest that SbRB initiate the assimilation of disaccharides in the wheat rhizosphere, which can then be utilized by a larger number of RhizCom members.

**Fig. 5.**
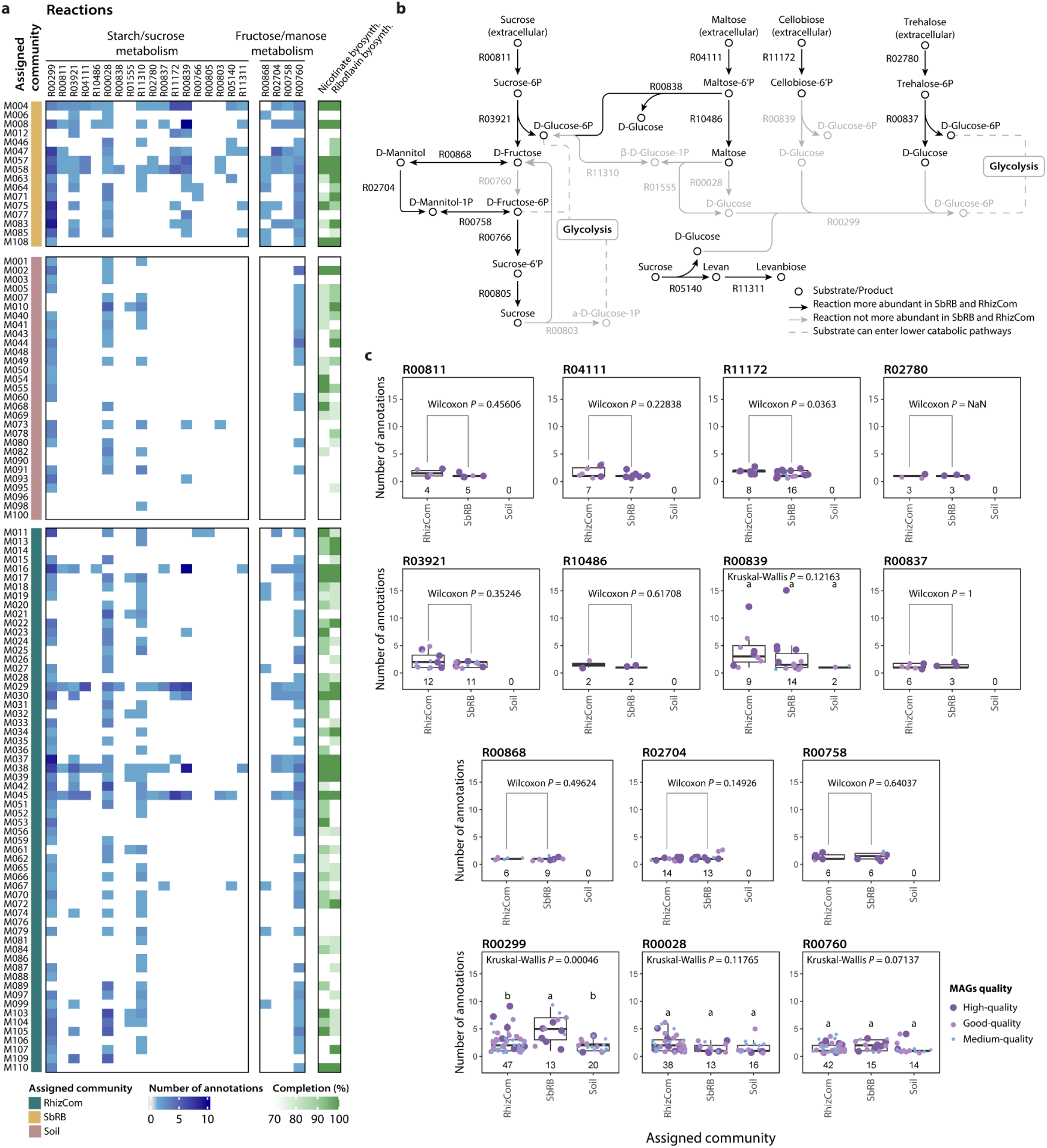
Metabolism of saccharides by SbRB drives the RhizCom assembly. **a**, Distribution of major KEGG reactions involved in specific saccharide metabolism within communities MAGs (blue) or the biosynthesis of nicotinate and riboflavin vitamins (green). Nicotinate: R07407,R00481+R04292+R03348+R03346+R02295. Riboflavin: R00425+R03459+R03458+ R07280+R07281+R04457+R00066+R00549+R00161. **b**, Reactions in pathways. Circles represent substrates and products metabolized by enzymes (arrows). Dotted lines represent substrates that can enter lower catabolic pathways (i.e. glycolysis). **c**, Number of annotations belonging to key KEGG reactions per MAG and assigned community. The numbers below the box plots indicate the number of MAGs that contain the reaction. Upper catabolic pathways of specific disaccharides (sucrose, maltose, cellobiose, and trehalose) are specific to SbRB/RhizCom members and not present in soil MAGs, while lower catabolic pathways (R0299, R000028, R00760) are present in most MAGs.

## Discussion

Crop health and productivity are highly dependent on the complex microbial communities they host^6,7,9^. While recent research highlights the superior role of native natural microbiomes over synthetic communities in supporting key plant functions^34^, limitations in obtaining reproducible natural plant-associated microbiomes challenge our ability to experimentally test microbiome manipulation strategies and gain insight into the fundamental principles governing microbial interactions and community dynamics.

In this work, we have demonstrated that sequential propagation of the wheat rhizobiome led to the recruitment of a stable, complex, and reproducible microbiome (Fig. 1). This is consistent with previous microbiome steady-states achieved after sequential growth in the rhizosphere of different host plants^20^, their phyllosphere^35^, or soil microbiomes^19^, suggesting robust environmentally mediated microbiome selection across hosts and habitats. The pronounced succession effect from the initial soil community to a converging rhizosphere microbiome (Fig. 1de) results from several host-mediated factors. Carbon sources exuded by young wheat roots^21,22^ primarily drove net community growth (Fig. 1f), favoring the proliferation of copiotrophs^36^. We used a 7-day time frame for each microbiome propagation cycle, representing a window of consistently high microbial activity that amplified the selective effect on community assembly. Additionally, plant immunity^37,38^, the secretion of plant secondary metabolites that influence bacterial assembly^24,25^, and the outcome of interbacterial interactions^15,16^ probably contributed to the assembly of the rhizosphere microbiome, which was exacerbated by successive microbiome propagation cycles. This favored the assembly of *Proteobacteria* and *Bacteroidetes* in the wheat roots to the detriment of soil-dwelling *Acidobacteria* or *Planctomycetes* (Fig. 1e). The resulting RhizCom was resilient to cryopreservation and regrowth on wheat roots, effectively maintaining a stable and reproducible rhizosphere community.

SbRB had a leading representation in the RhizCom after coalescing with members of the soil microbiome, most of which were acquired during the multiple cycles of community succession (Fig. 2, Supplementary Table 3). Although the SbRB community was based on a limited set of plants during a single propagation cycle, it was sufficient to capture 50% of the relative abundance of the RhizCom. Moreover, integrating the information from the succession cycles, which accounted for a total of 96 individual plants, 91% of the relative abundance of RhizCom was seed-borne. The dominance of seed-transmitted bacteria in the rhizosphere microbiome of various young plants has been observed previously^28^, and likely reflects instances of priority effect where pioneer members of the seed microbiota strongly shape the root-associated community^39,40^. We argue that the rapid habitat changes during early stages of plant development, and the periodic resetting to the initial habitat state, further stressed primary succession. This effect would have strongly favored the assembly of bacteria adapted to the rapid utilization of carbon exuded by developing roots and may have limited the establishment of late-arriving soil bacteria, e.g., by niche preemption^39^, although this could still occur at later stages of habitat development (Fig. 6). Nevertheless, 8% of the relative abundance of the RhizCom was derived from the soil community. This could be explained by some members having similar metabolic capabilities to SbRB, strong competitive abilities, or be evidence for instances of niche facilitation^41^. Whether these processes will maintain the reproducibility of the RhizCom in later stages of habitat development^42^ remains to be investigated.

**Fig. 6.**
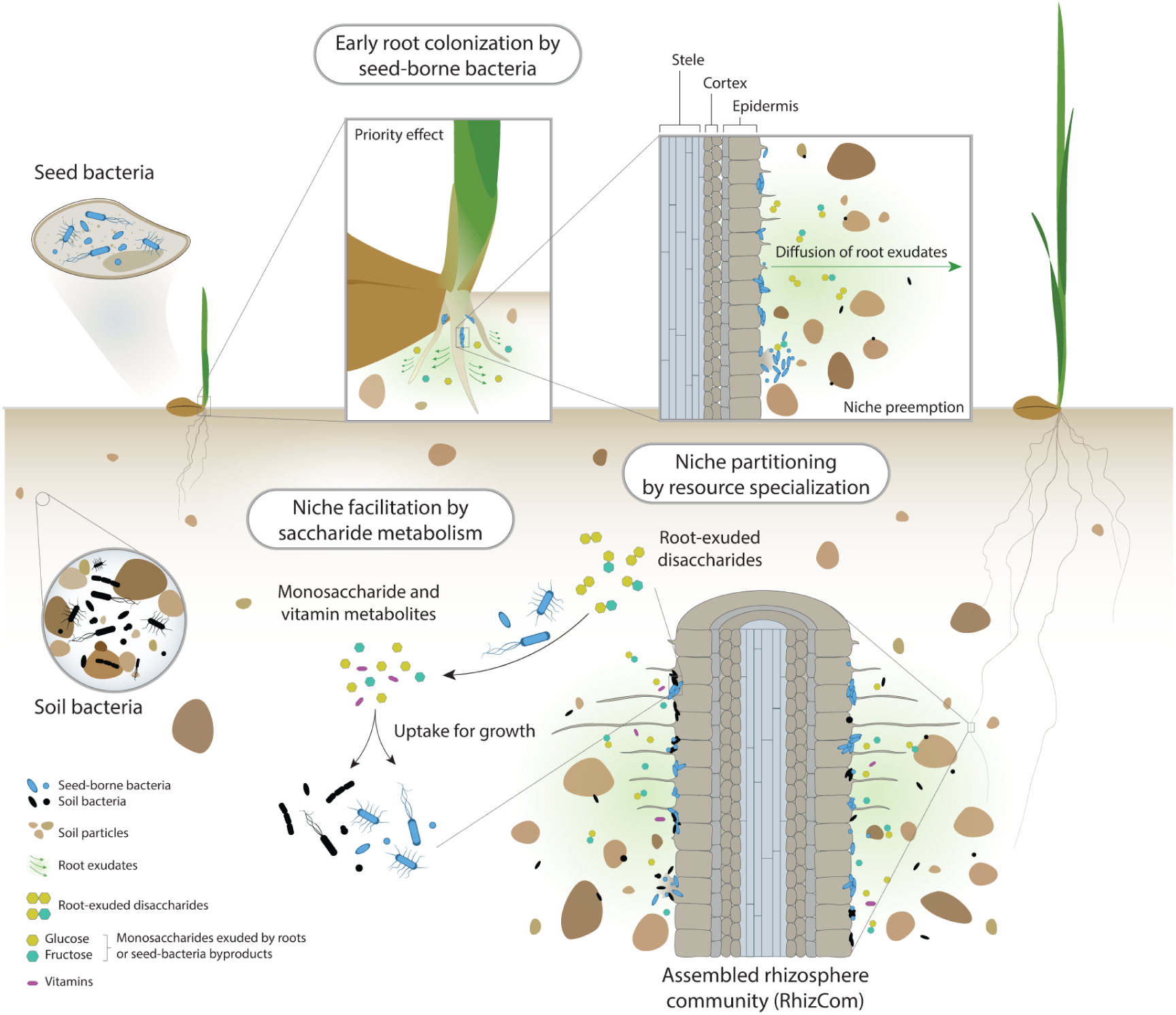
Priority effects of heritable seed-borne bacteria drive early assembly of the wheat rhizosphere microbiome. Representation of the processes that occur during early assembly of the wheat rhizosphere microbiome. Upon germination, seed bacteria are the first to colonize root surfaces, using root exudates as carbon sources. This early colonization leads to the depletion of readily available nutrients in the rhizosphere (niche preemption), preventing early colonization by soil-dwelling bacteria. In later stages of plant development, the specialized metabolism of root-exuded disaccharides by seed-borne bacteria allows them to persist when competition for primary resources from late-arriving soil bacteria increases through niche partitioning. The resulting accumulation of disaccharide metabolism byproducts, glucose and fructose, by SbRB facilitates the colonization of the wheat rhizosphere by both soil and seed-borne bacteria. In addition, the biosynthesis of the vitamins nicotinate and riboflavin by SbRB supports the establishment of auxotrophic bacteria, promoting community coalescence and increasing the complexity of the rhizosphere microbiome.

The patterns of functional specialization observed among the communities (Fig. 3) were related to the exploitation of their different habitats. In the soil community, enriched functions characteristic of oligotrophic bacteria^43,44^ such as *Actinobacteria* or *Planctomycetes*, reflected their broad energy metabolism and secondary metabolite production (Fig. 3b, Extended Data Fig. 4, Supplementary Fig. 7), supporting survival in resource-limited, variable environments^44–46^. This broad metabolic capacity was further highlighted by the higher metabolic diversity of the soil community compared to the RhizCom (Fig. 3e). In contrast, the RhizCom exhibited a functional profile dominated by enrichment in transcription, signal transduction, and other traits characteristic of metabolically active copiotrophic bacteria^36,47^. Additionally, traits required for successful colonization and resource competition in the rhizosphere were enriched in the RhizCom, including biofilm formation, motility, or carbohydrate and amino acid metabolism^48–50^ (Supplementary Fig. 7). The biosynthesis of extracellular polymeric levan may have additionally supported the effective colonization of the rhizosphere^51^ by SbRB (Fig. 5).

In contrast to the other two communities, SbRB did not show specific enrichment in any of the broad functional categories, except for membrane transport (Fig. 3ab). This likely reflects that the functional traits of SbRB were largely integrated into RhizCom during sequential community succession. However, specific enriched functions in SbRB were related to carbohydrate metabolism, such as sugar transport and metabolism systems (*msmEFG, bglAF, fruAB,* Fig. 3d), and, together with the RhizCom, also iron acquisition. The enrichment of these functions in SbRB members corresponds to essential traits in early instances of habitat exploitation by the seed microbiota, including the utilization of saccharides exuded by developing plant roots^50,52^ and scavenging strategies to secure limiting iron^53,54^. Indeed, the resource specialization of SbRB in the metabolism of sucrose, maltose, cellobiose and trehalose (Fig. 5), disaccharides commonly exuded by plant roots or produced during cellulose degradation^50^, emphasizes their role as early metabolic specialists, particularly *Pantoea*, *Paenibacillus* and *Priestia*. Since disaccharides are less readily available carbon sources compared to glucose and fructose, which are also exuded by wheat roots^55,56^, SbRB may have initially focused on the exclusive use of monosaccharides in the absence of early soil-dwelling competitors due to priority effects, leading to the preemption of simple sugars^39^. As competition for primary resources increases with the arrival of later-arriving bacteria, specialized disaccharide metabolism by established SbRB would have allowed them to persist in the rhizosphere through niche partitioning. The monosaccharide byproducts of this metabolism could then accumulate, in turn facilitating broader bacterial colonization, suggesting a secondary role for niche facilitation (Fig. 6). This sequence of niche partitioning and facilitation by saccharide metabolism allows SbRB to maintain their dominance in the rhizosphere at later stages of plant development. By producing the vitamins nicotinate and riboflavin (Fig. 5a), SbRB may have further contributed to niche construction, enabling both soil bacteria and other seed-borne members lacking these functions to coalesce into the RhizCom^57^ (Fig. 6). While this study highlights the key role of SbRB in shaping early rhizobiome assembly, future work should experimentally resolve the contributions of niche facilitation during community succession.

Our work demonstrates that sequential propagation is a key strategy for achieving reproducible rhizosphere microbiomes, driven by strong host-mediated effects under constrained habitat conditions. Priority effects and niche facilitation through saccharide metabolism of seed-borne bacteria were key determinants for the assembly of the rhizosphere community, highlighting the processes that drive primary succession during early habitat development. These results contribute to our understanding of microbial community dynamics and provide valuable strategies for experimentally testing microbiome manipulation approaches aimed at improving crop productivity and health.

## Methods

### Initial soil sampling and characterization

The soil used for the wheat rhizosphere microbiome recruitment was collected from a mixed plant species grassland bordering a wheat field in the region of Grandcour, Switzerland (46.884947N, 6.922562E) in April 2021, as previously described^9^. The bacterial load of the soil was determined following mixing soil and mineral medium^18^ (MM, 1:1). The mixture was incubated for 1 h at room temperature (RT) with shaking (180 rpm) and then centrifuged at 300 × *g* for 1 min to pellet soil debris. The supernatant was further centrifuged at 4,500 × *g* for 5 min. The resulting pellet was washed twice with MM. Serial dilutions were plated onto R2A medium and incubated at 25 °C. Colony forming units (c.f.u.) were enumerated after 48 h. Three independent samples and 10 technical replicates per sample were performed, resulting in an average of 1.18 × 10^6^ ± 3.43 × 10^5^ c.f.u. g^-1^ of soil.

### Sequential propagation of the wheat rhizosphere microbiome

To obtain a reproducible wheat rhizosphere microbiome, we used a previously designed microcosm^18^. Each microcosm contained a soil matrix (SM), soil extract (SE), a bacterial inoculum, and a pre-germinated 2-day-old wheat *Triticum aestivum* cv. Arina) seedling (Extended Data Fig. 1). The obtention of the SM, SE and the germination of the wheat seeds is described in the Supplementary information. The bacterial inoculum was either a soil wash (SW) containing the bacterial suspension extracted from the soil, or a rhizosphere wash (RW) obtained after each propagation cycle (see Supplementary information for details). The microcosms were maintained in a Percival PGC-7L2 plant growth chamber at 22/18 °C, 16/8 h light/dark photoperiod (160 μE m^-2^ s^-1^ light intensity), and a relative humidity of 70% for seven days. Sixteen tubes were prepared per propagation cycle. After seven days, the RW of pools of four plants was recovered and used for c.f.u. enumeration of bacteria, total DNA extraction, and for the reinoculation of the next cycle. For c.f.u. enumeration of bacteria, aliquots of the RW were serially diluted in MM and plated on R2A plates. Eight technical replicates per pool of each four plants were used for c.f.u. enumeration. For DNA extraction, 10 mL of the RW was centrifuged at 7,000 × *g* for 15 min to pellet bacterial cells, which were cryopreserved at −20 °C until DNA extraction (see below). Finally, another 10 mL of the four RW of all four rhizosphere pools was combined into a single suspension, which was centrifuged at 7,000 × *g* and the supernatant discarded. The pellet was resuspended in 40 mL of SE, and 2 mL of the suspension was inoculated per tube for the next propagation cycle. A total of six propagation cycles were performed, after which the final RW was used for c.f.u. enumeration of bacteria and total DNA extraction as described above, and to preserve the microbial community by storing the suspension at −80 °C in glycerol 50% (v/v).

The cryopreserved wheat rhizosphere community stored at −80 °C was recovered by thawing and centrifuging two aliquots at 7,500 × *g* for 1 min to pellet the cells, and then washing the pellet twice with MM. Finally, the pellet was resuspended in 40 mL of SE which served to inoculate 16 microcosms with wheat seedlings, as described. To recover the seed-borne rhizosphere bacteria (SbRB), microcosms were set up for a cycle (seven days) of plant growth, following the same protocol as described above, without the addition of an inoculum.

### DNA extraction, library preparation, and sequencing

Total DNA from samples was extracted using the DNAeasy PowerSoil Pro kit (Qiagen), following the manufacturer specifications. DNA concentrations were measured using the Qubit dsDNA HS assay kit (Invitrogen). Libraries of the 16S rRNA gene were prepared using 10 ng of extracted DNA per PCR reaction, using primers targeting the 16S rRNA V3-V4 region following the Illumina 16S Metagenomic Sequencing Library protocol, as previously described^18^. The A Nextera XT index kit (v2, Illumina) was used for indexing. Samples were then quantified and pooled in equal amounts for sequencing. Samples were spiked with 25% PhiX control DNA and paired-end sequenced using an Illumina MiSeq v3 instrument running for 600 cycles. For shotgun metagenome sequencing, samples from the initial soil, after the sixth reinoculation cycle, and SbRB were selected. A total of 100 ng of DNA per sample were used for library preparation, using the Nextera DNA Flex library protocol (Illumina). Metagenome samples were paired-end sequenced using a NovaSeq 6000 instrument running for 300 cycles and using S2 flow cells. Samples were sequenced at the Lausanne Genomic Technologies Facility (Lausanne, Switzerland). The raw reads from both, 16S rRNA gene amplicons and shotgun metagenome were quality filtered and trimmed with fastp^58^ v0.23.2, using the adaptor autodetection option.

### 16S rRNA gene amplicon analyses

Quality-filtered 16S rRNA gene amplicon reads were analyzed following the DADA2^59^ v.1.16.2 pipeline, as previously described^18^. SILVA database^60^ v138.1 was used for taxonomy assignation at 99% sequence identity. ASV sequences were used to construct a maximum-likelihood phylogeny as previously described^18^. Data was imported into phyloseq^61^ v1.42.0 R package. ASVs classified as mitochondria, chloroplasts or with prevalence < 0.05 across samples were removed. Cumulative sum scaling (CSS) normalization^62^ was applied to ASV counts. Relative abundances at the bacterial class taxonomic rank were analyzed and represented within the 500 most abundant ASVs, using the *plot_bar* function within phyloseq R package. Observed ASVs and the Shannon diversity index were calculated and plotted with the *plot_richness* phyloseq R function. Phylogenetic diversity (Faith’s PD) was calculated using the *pd* function within the picante v1.8.2 R package^63^. A principal coordinate analysis (PCoA) was performed with the *ordinate* phyloseq R function and using Bray-Curtis dissimilarities. Bray-Curtis dissimilarities were also used for a complete-linkage hierarchical clustering using the ComplexHeatmap^64^ v2.9.2. R package. Differential abundance analysis of ASVs across samples was determined using DESeq2^65^ v1.38.0 R package as previously described^18^. The phylogenies of ASVs were visualized using the ggtree^66^ v3.6.0 and ggtreeExtra^67^ v1.8.1 R packages.

### Metagenome analyses

Quality-filtered reads were mapped against a reference wheat genome (NCBI GenBank acc. no. GCA_903993985.1) to remove host contamination, using Bowtie2^68^ v2.5.1. SAMtools^69^ v.1.7 was used to retrieve unmapped reads. To assess sample coverage, Nonpareil3^70^ v3.401 was used with the host-removed reads, using a *k*-mer of 31 and 100,000 random reads. Reads were downsampled to a maximum of 0.6 billion reads per replicate to account for computational limitations, and subsequently normalized by *k*-mer coverage using BBnorm v37.62 (sourceforge.net/projects/bbmap). A first normalization run was performed using a target *k*-mer coverage of 50 per replicate paired-end reads. Reads from the sample replicates were then pooled, and a second normalization run also using a target *k*-mer coverage of 50 was performed. To reduce the complexity of the soil sample, pooled reads with a *k*-mer coverage below 15 were discarded. The number of reads obtained at each step was assessed with Seqkit^71^ v2.2.0 as summarized in the Extended Data Fig. 2. Normalized pooled reads from the last reinoculation cycle (RhizCom), SbRB, and soil were co-assembled using MEGAHIT^72^ v1.2.9 with the options *--presets meta-large* and *--kmin-1pass*, and a minimum contig length of 1,000 bp. The assemblies were analyzed using QUAST^73^ v5.2.0, resulting in 1.54 million contigs, totaling 5.2 billion bp (Extended Data Fig. 2).

The resulting contigs were analyzed using the SqueezeMeta^74^ v1.6.2 pipeline. Briefly, open reading frames (ORFs) were predicted using Prodigal^75^, followed by similarity searchers using the GenBank non-redundant (nr)^76^, eggNOG^77^, and KEGG^78^ databases with diamond^79^ v2.0.15.153. Gene searches were also performed using HMMER3^80^ against the Pfam^81^ database. The databases were downloaded in May 2023. The least common ancestor (LCA) algorithm^74,82^ was applied to assign genes at different taxonomical ranks using the search results against the GenBank nr database. Reads after host removal were used for mapping against contigs using Bowtie2^68^ v2.5.1 and to calculate coverage and abundance estimations (normalized transcripts per kilobase million, TPM) for genes. Results were analyzed using the SQMtools^83^ v1.6.2 R package.

For differential abundance analyses, we defined a minimum abundance threshold ≥ 1,000 across samples to be included in the analysis (Supplementary Fig. 8), using DESeq2 as specified above. KEGG annotations were represented in a ternary plot using the ggtern^84^ v3.5.0 package. Mean TPMs per sample were used to calculate the ternary coordinates per KEGG annotation, with a minimum TPM threshold ≥ 0.5 across samples. Differential abundance results were used to categorize a KEGG annotation as enriched if the KEGG annotation was significantly more abundant in a sample in all comparisons that included that sample, or as suppressed if the annotation was not significantly more abundant in any of the comparisons. Intermediate cases and those where the annotation was not significantly enriched in two or three comparisons, were considered as balanced.

### Analyses of MAGs

Contigs were binned into MAGs using CONCOCT^85^ v1.1.0, MetaBAT 2^86^ v1:2.15, and MaxBin^87^ v2.2.6. Resulting bins were dereplicated and aggregated into a single set of MAGs using DASTool^88^ v1.1.1. The coverage of MAGs across samples was estimated by mapping host-removed reads to bins, as described above. MAGs statistics were calculated using CheckM^89^ v1.1.6, and taxonomic assignment was based on the bin gene consensus from SqueezeMeta^74^. The quality of MAGs was assessed according to standard metrics^90^. The maximum-likelihood (ML) phylogeny of 110 high-to-medium quality MAGs was built with PhyloPhlAn^91^ v3.0.67, using its database of 400 universal amino acid marker sequences^92^, and a *--diversity medium* option. Bootstrap support was assessed using RAxML^93^ v8.2.12, with 1,000 rapid bootstrap inferences followed by a thorough ML search for the best scoring tree. Results were imported into R and plotted using the ggtree and ggtreeExtra R packages. Seven MAGs without sufficient PhyloPhlAn marker sequences to be included in the phylogeny were discarded. A MAG was assigned to a community if its mean TPM percentage was ≥ 1.5x that of the other two communities. If this condition was not met, the MAG was assigned to communities where the mean TPM percentage difference was < 1.5x.

### Statistical analysis

Data used for statistical analyses were tested using the Saphiro-Wilk normality test, and the Levene’s test for homogeneity of variances. Data that did not have a normal distribution or homogeneous variances were analyzed with the agricolae^94^ v1.4-5 R package, using the non-parametric Kruskal-Wallis rank sum test, with Fisher’s least significant difference (LSD) post hoc criterium, and correction of *P* values using the false discovery rate (fdr). Pairwise comparisons were performed using the Wilcoxon rank-sum test within the stats v4.4.1 R package. Data that followed a normal distribution and with homogeneous variances were analyzed using PERMANOVA (permutational multivariate analyses of variance) or ANOSIM (analysis of similarity), using with the adonis2 or anosim functions, respectively, within the vegan^95^ v1.5-4 R package, and using 9999 permutations. Spearman correlations were calculated using the *stat_cor* function within the ggpubr R package, and data was fitted to a general additive model (GAM), using a *k* = 3 with the *geom_smooth* ggplot2 v3.3.6 R function, and a confidence interval of 0.95.

## Data availability

Raw reads from both 16S rRNA gene amplicons and metagenomic shotgun have been deposited in the NCBI Sequence Read Archive (RSA) database and are publicly available under the BioProject accession number PRJNA1169405. Other raw data generated in this study are provided in the Supplementary information or in the GitHub repository https://github.com/dgarrs/RhizCom.

## Code availability

The code used for the analysis of the 16S rRNA gene amplicon data, metagenomic shotgun data, and other analyses reported in this study is available on GitHub (https://github.com/dgarrs/RhizCom).

## Acknowledgements

We thank Julia Vorholt and Alan Pacheco at ETH Zurich for useful discussions and feedback on the manuscript. We are also grateful to Jordan Vacheron for his input during research discussions. We thank Caterina Matasci (Delley seeds and plants Ltd, Switzerland), for providing us with the wheat seeds used in this study. We also thank the Lausanne Genomic Technologies Facility in Lausanne, Switzerland for the sequencing service. This work was supported as a part of the NCCR Microbiomes, a National Centre of Competence in Research, funded by the Swiss National Science Foundation (grant numbers 180575 and 225148). DG-S was funded by the Fondation pour l’Université de Lausanne.

## Author contributions

Conceived and designed the experiments: DGS and CK.

Preformed the experiments: DGS.

Analyzed the data: DGS.

Contributed materials/analysis tools: DGS.

Wrote the paper: DGS and CK.

## Competing interests

The authors declare no competing interests.

## Extended Data

**Extended Data Fig. 1.**
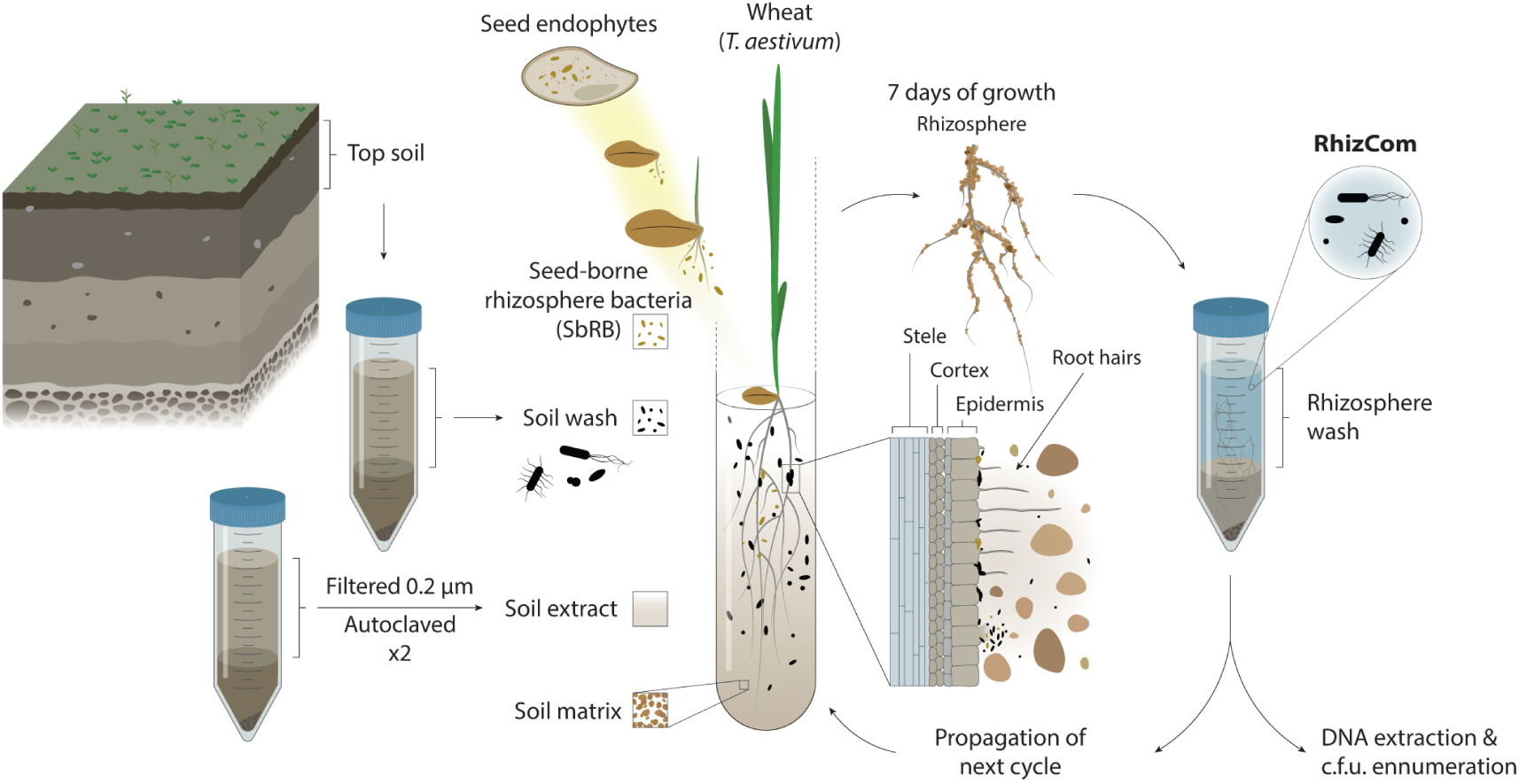
Schematic representation of the experimental design and microcosm used in this study. The top bulk soil from a Swiss grassland bordering a wheat field was used to obtain a soil wash containing the soil microbial suspension, and a filtered and sterile soil extract to supplement the microcosm with soluble micronutrients and molecules present in the soil. Both the soil wash and the soil extract were introduced into sterile microcosms consisting of glass tubes containing a soil matrix (0.5-4 mm silt). A two-day-old wheat (*Triticum aestivum* var. Arina) seedling was then planted. After seven days of plant growth and microbiome development, the rhizospheres (roots and attached soil matrix) were collected, and the microbial cells were washed. The resulting rhizosphere wash was used to reinoculate the next propagation cycle, to extract total DNA for 16S rRNA amplicons and metagenome shotgun sequencing, and to enumerate c.f.u. of bacteria. The rhizosphere wash obtained at the end of the sixth propagation cycle contained the microbial community considered as the replicable rhizosphere community (RhizCom). Seed endophytes that were able to leave the seed tissues and establish in the wheat rhizosphere in the absence of the soil wash inoculum, were considered the seed-borne rhizosphere bacteria (SbRB).

**Extended Data Fig. 2.**
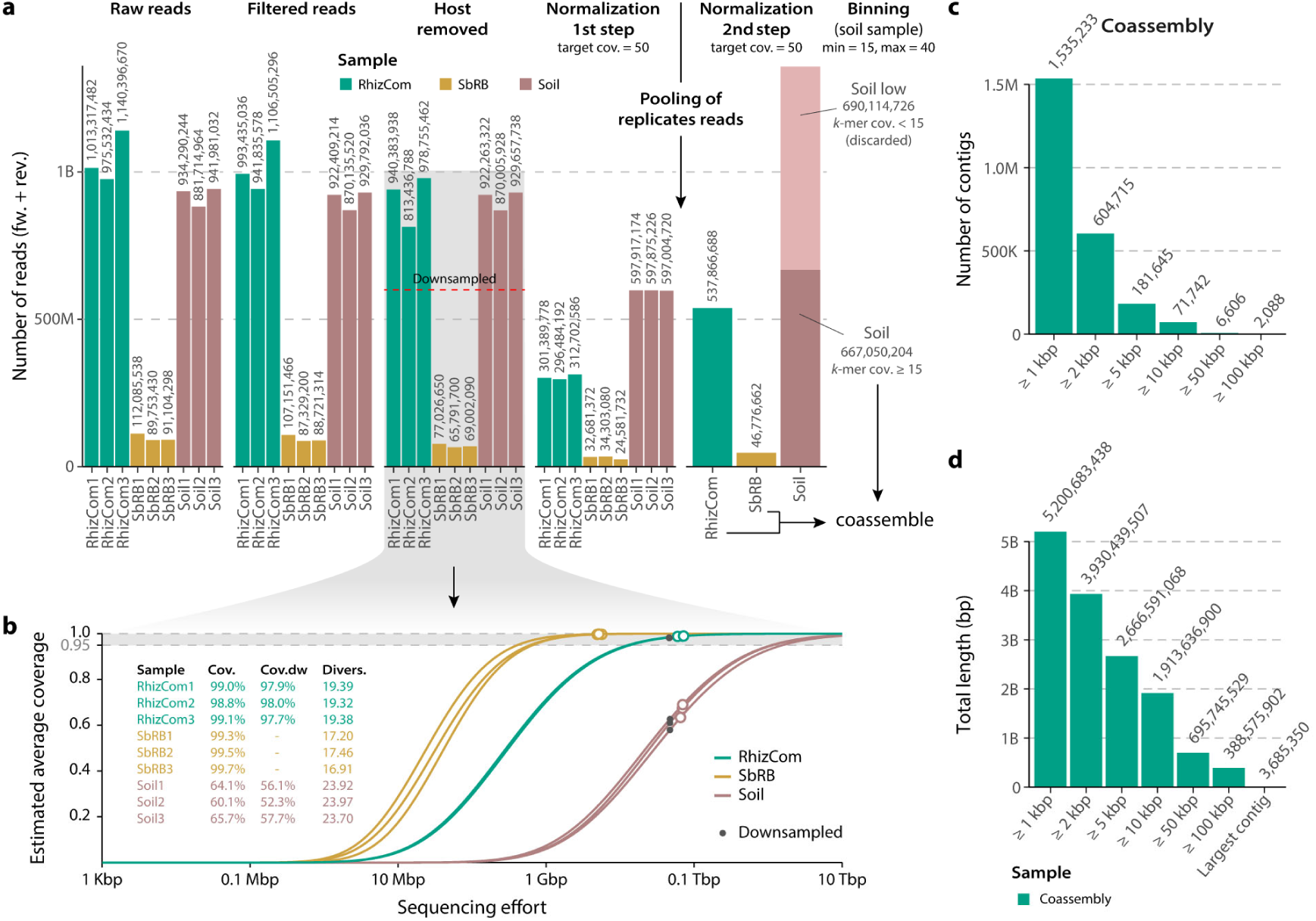
Processing of metagenome sequencing reads and coassembly statistics. **a**, Combined number of forward and reverse reads obtained at each processing step in the RhizCom, SbRB and soil replicates. After host removal of host reads, reads were downsampled to a maximum of 600 million reads per sample (red dashed line) to account for computational limitations. **b**, Estimated sequence coverage per sample based on host-removed reads as a function of the sequencing effort. Empty dots along the curves indicate the average coverage achieved per replicate. Full black dots indicate coverage obtained after downsampling. Values of sample coverage (Cov.), sample coverage after downsampling (Cov.dw) and sample sequence diversity (Divers.) are indicated. **c,d**, Number of contigs (**c**), or total length in base pairs (bp, **d**) after the coassembly of samples.

**Extended Data Fig. 3.**
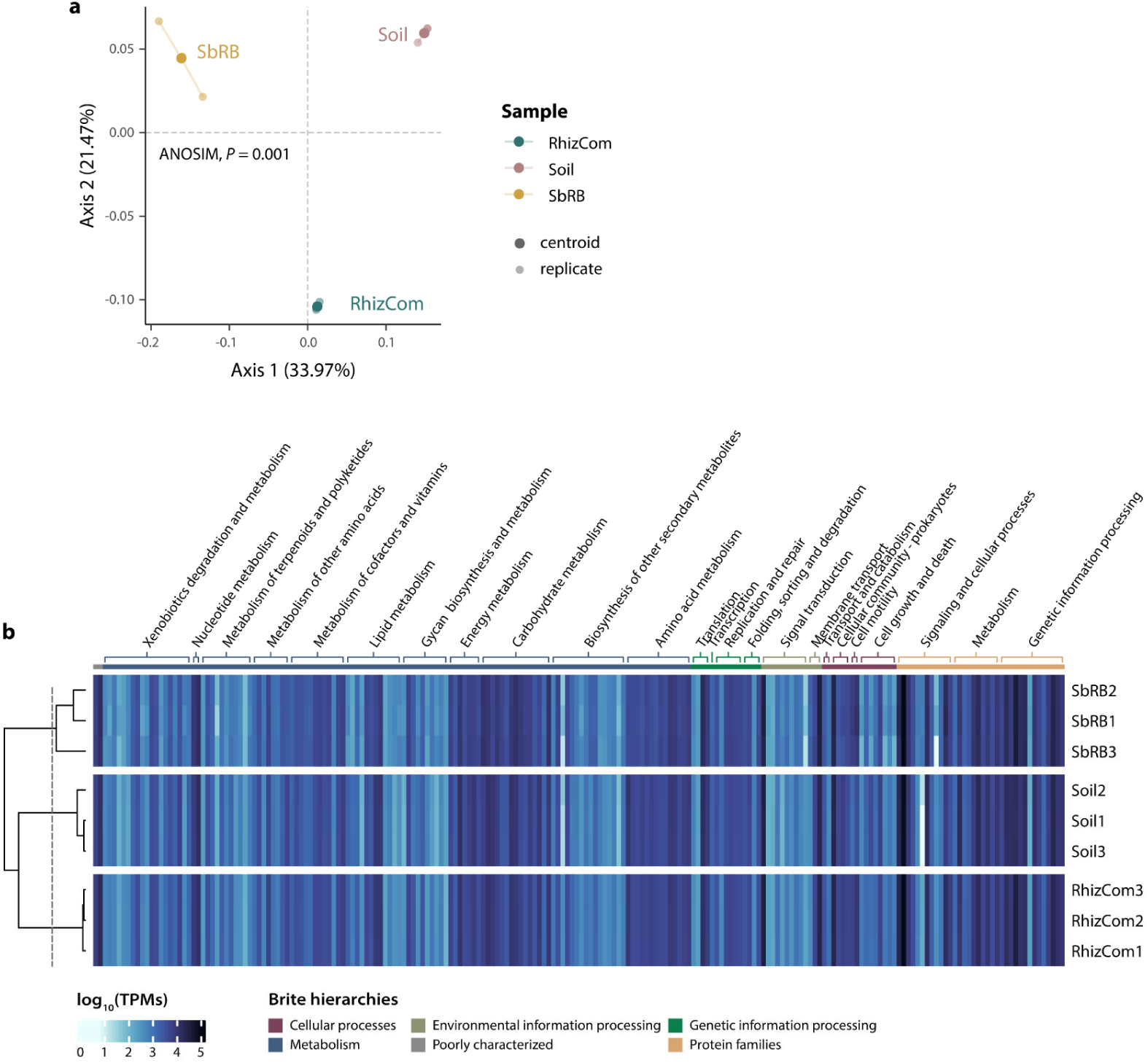
Overall functional content of samples. **a**, Principal coordinate analysis (PCoA) of samples based on KEGGs TPM showing the two first axes. Statistical differences between samples were assessed based on Bray-Curtis dissimilarities using ANOSIM (Analysis of Similarities), using 9999 permutations. **b**, Distribution of KEGG BRITE hierarchies across samples. Samples were clustered based on the complete linkage of Euclidean distances. Only hierarchies with a total sum > 1000 TPMs across samples were represented.

**Extended Data Fig. 4.**
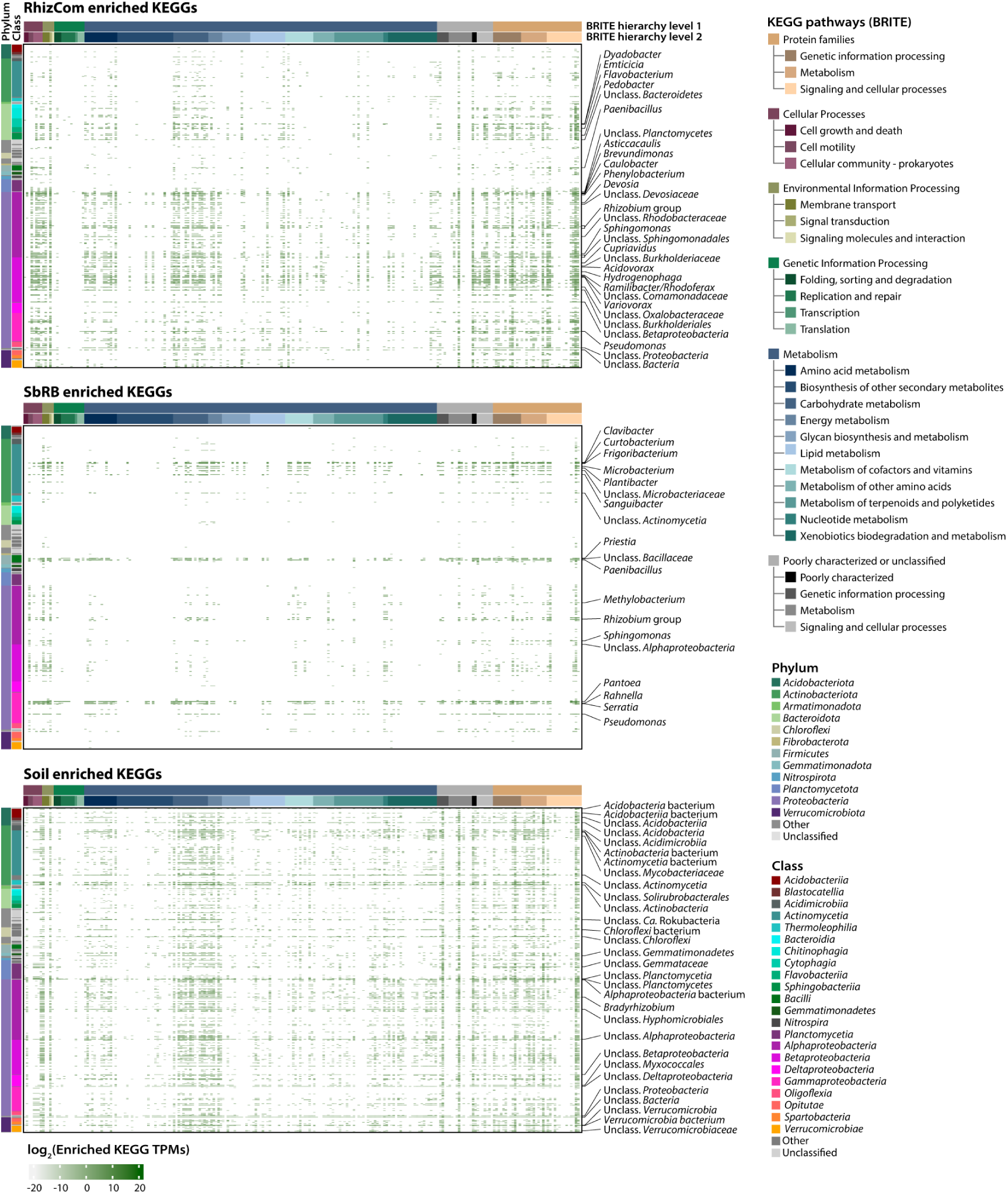
Taxonomic distribution of enriched KEGG annotations across communities. Enriched KEGG annotations across communities based on DESeq2 differential abundance comparisons of TPMs. TPMs were summed per KEGG annotation and taxonomic rank at the genus level.

## Supplementary Datasets

**Supplementary Table 1.** Enumeration of colony-forming units of the rhizosphere microbiome during the propagation cycles.

**Supplementary Table 2.** Number of taxa per sample obtained by 16S rRNA profiling.

**Supplementary Table 3**. Contribution of soil wash and SbRB to the RhizCom.

**Supplementary Table 4.** Metagenome analysis statistics.

**Supplementary Table 5.** Kruskal-Wallis rank sum test of KEGG annotations TPMs across categories.

**Supplementary Table 6.** KEGG annotations per samples dominance categories. **Supplementary Table 7.** Differential abundance analysis on KEGG annotations. **Supplementary Table 8**. Characteristics of high-to-medium-quality MAGs.

## Supplementary methods

### Microcosm and experimental design

Each microcosm^1^ consisted of a sterile 3.3 borosilicate glass tube (diameter Ø25 mm x 200 mm, Fisher Scientific) closed with sterile Magenta 2-way polypropylene caps (Sigma-Aldrich). Each microcosm contained a soil matrix (SM), soil extract (SE), a bacterial inoculum, and a pre-germinated 2-day-old wheat (*Triticum aestivum* cv. Arina) seedling. The SM, which served as a substrate for plant root and microbiome development, comprised 20 g of 0.5 – 4 mm silt that had been sieved and autoclaved twice (120 °C, 15 min). The SE, which provided micronutrients and minerals from the soil environment, was prepared by mixing soil with mineral medium^1^ (MM, 1:1 ratio) for 1 h at 180 rpm, followed by centrifugation at 4,500 × g for 15 min. The resulting supernatant was filtered through 0.2 µm cellulose acetate filters (Dutscher) and autoclaved twice, and its final pH was 6.48 ± 0.03. The bacterial inoculum was either an initial soil wash (SW) containing a bacterial suspension extracted from the soil, or a rhizosphere wash (RW) obtained after each reinoculation cycle (see below). The SW was prepared by mixing soil with MM (1:1 ratio) for 1 h at 180 rpm, followed by a low-speed centrifugation at 300 × g for 1 min to remove soil debris. The supernatant was then centrifuged at 4,500 × g for 15 min, and the resulting pellet washed twice with MM. The pellet was resuspended in SE and considered as the soil wash (SW) suspension containing cells from soil. Two mL of the SW was added to the 20 g of SM in each glass tube. Finally, a pregerminated wheat seedling was placed in each tube. For pre-germination, seeds were surface disinfected in 4% NaClO for 15 min, thoroughly washed with sterile distilled water, and germinated on 0.85% (w/v) agar plates in the dark at 22 °C for 2 days. The microcosms were incubated in a Percival PGC-7L2 plant growth chamber at 22/18 °C, 16/8 h light/dark photoperiod (light intensity, 160 μE m^-2^ s^-1^), and 70% relative humidity. After seven days of plant growth in the microcosm, the plants were removed from the tubes and their roots were gently shaken to detach any loosely adhering particles of SM. Roots with the adhering SM were cut off at the point of emergence from the seed, representing the rhizosphere, which also encompasses the rhizoplane. The rhizospheres of four plants were pooled and weighed. Twenty-five mL of MM was added to each sample pool, followed by vortexing for 20 min. The samples were then centrifuged at 300 × *g* for 1 min to pellet soil matrix debris, and the resulting supernatants were considered the rhizosphere wash (RW), which was used for total DNA extraction, c.f.u. enumeration, and prepare the reinoculation solution. The reinoculation solution was prepared by mixing 10 mL of the RW of four replicates (total of 40 mL of RW), which was then centrifuged at 4,500 × *g* for 5 min. The supernatant was discarded, and the pellet was resuspended in 40 mL of SE. This solution was used to inoculate the next cycle.

## Supplementary figures

**Supplementary Fig. 1.**
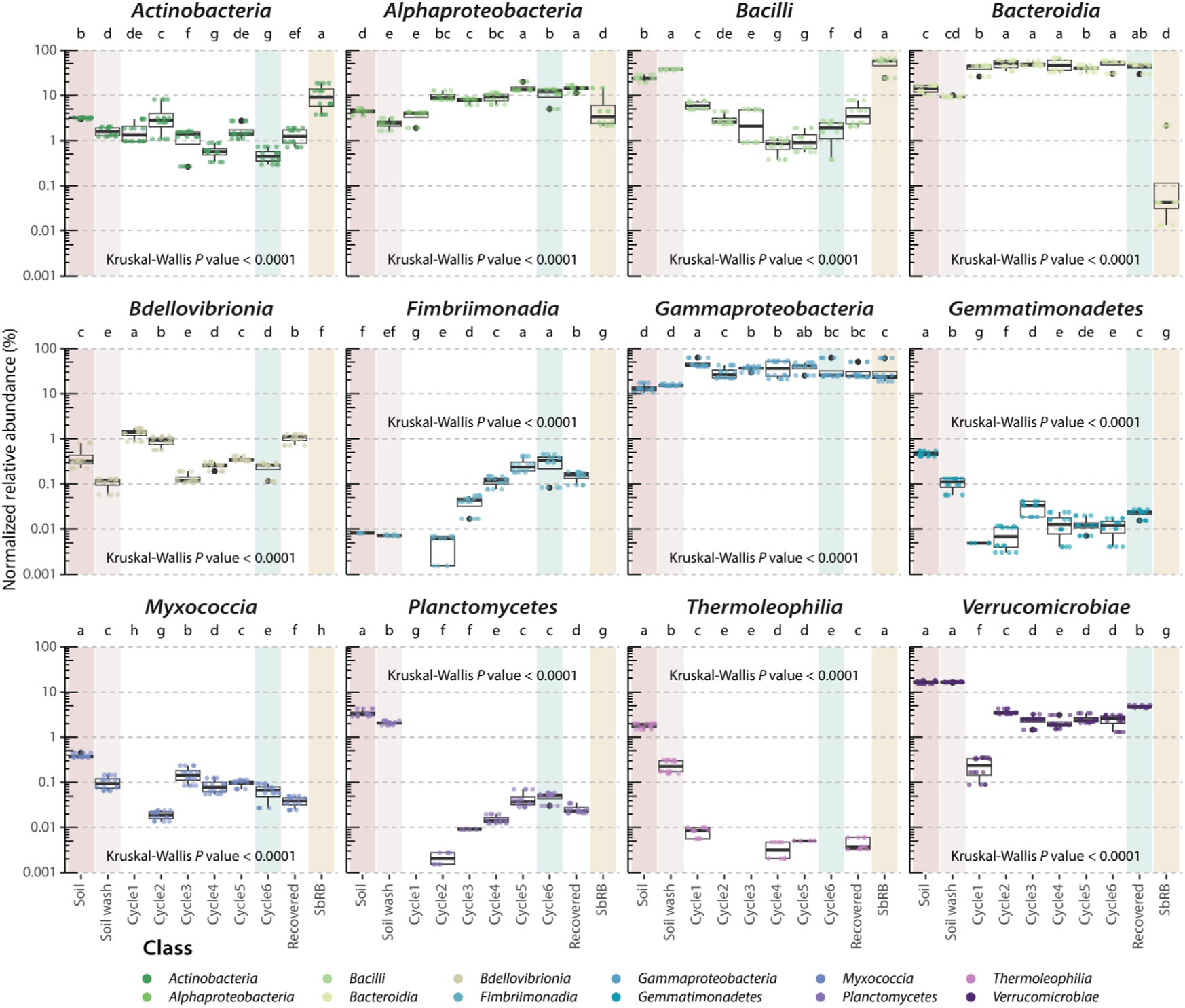
Differences in relative abundance of bacterial classes across samples. CSS-normalized relative abundance of the top bacterial classes across samples. Letters denote different statistically significant groups (*P* value < 0.05) using Kruskal-Wallis rank sum test with LSD post hoc analysis and *P* value corrected by fdr.

**Supplementary Fig. 2.**
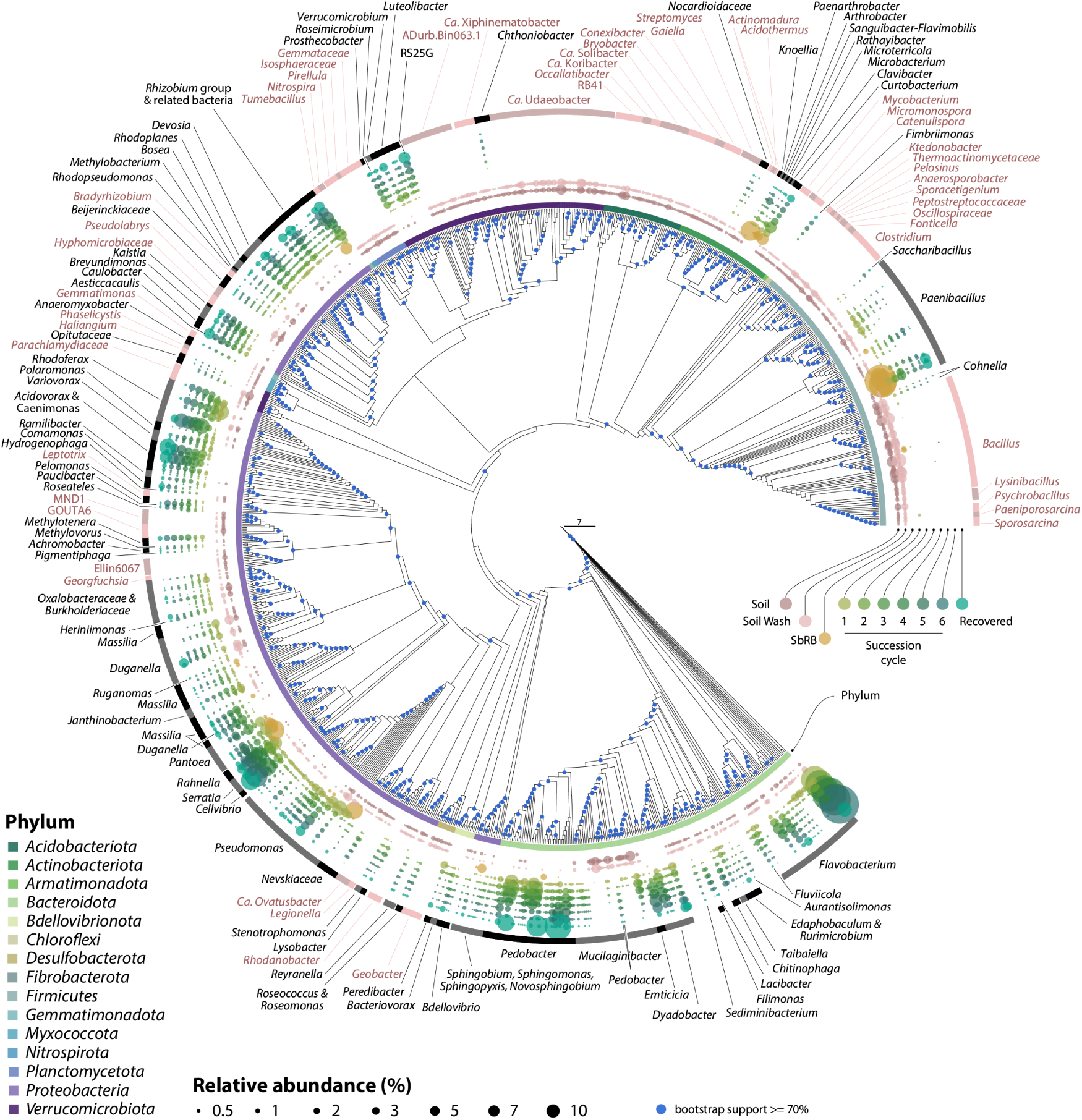
Selection of ASVs during the succession of the wheat rhizobiome. The phylogenetic tree was built using ASVs with a total relative abundance ≥ 0.005%. Names of relevant genera selected (black/grey) or not (pink/dark pink) after the succession are indicated. From inwards to outwards, colored dots represent ASVs, with sizes corresponding to their relative abundance in soil, soil wash, SbRB, succession cycles and the recovered rhizosphere communities.

**Supplementary Fig. 3.**
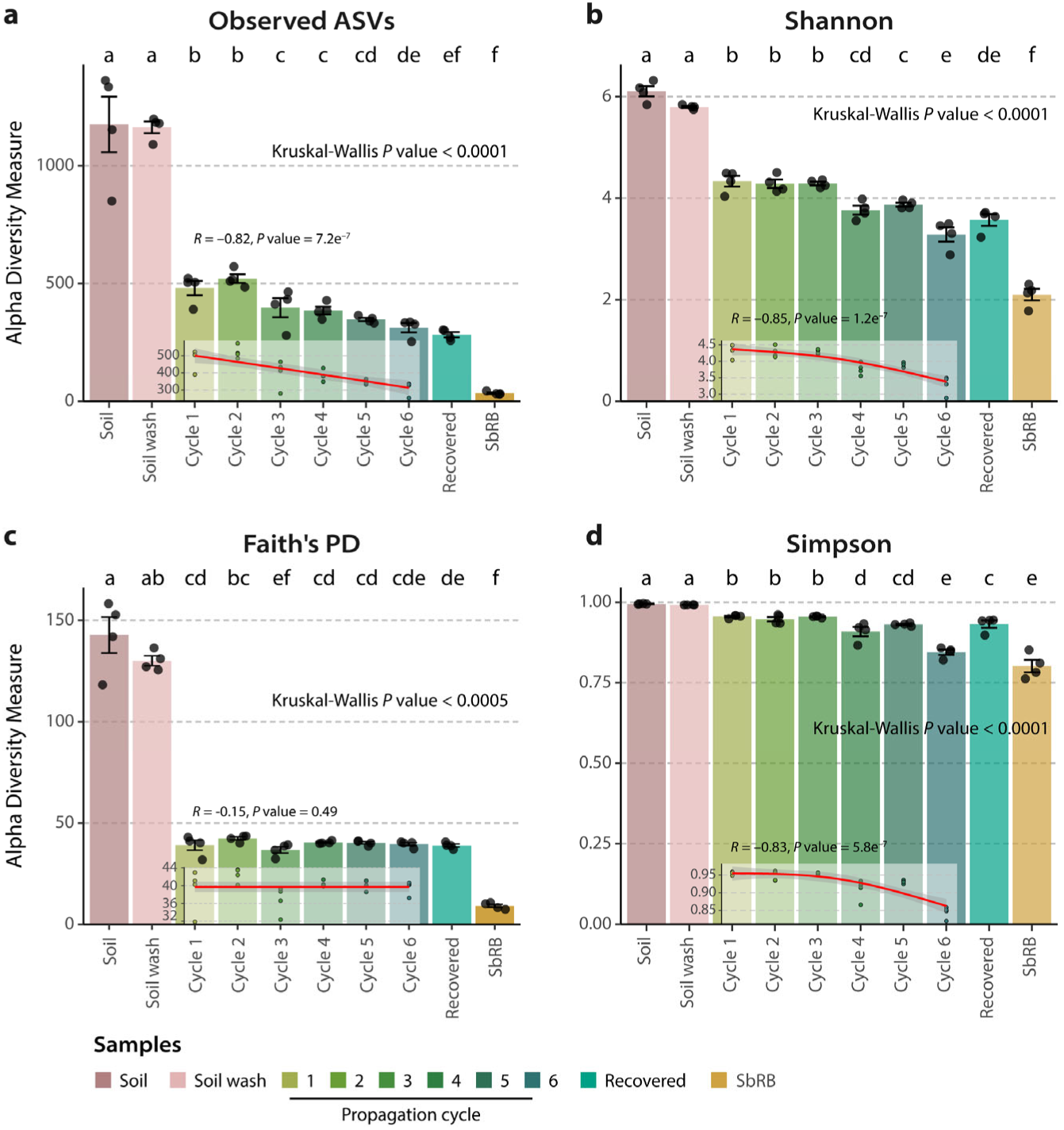
Alpha diversity of samples across succession cycles. Number of observed ASVs (**a**), Shannon diversity (**b**), Faith’s phylogenetic diversity (PD, **c**), and Simpson diversity (**d**) across samples are indicated as bars (mean values) with the standard error. Black dots indicate values of individual replicates. Letters denote different statistically significant groups (*P* value < 0.05) using Kruskal-Wallis rank sum test with LSD post hoc analysis and *P* value corrected by fdr. Spearman correlation of each alpha diversity measure and succession cycle is shown within the plots. Correlation coefficient (*R*) and *P* value are indicated. Curves represent the general additive model fit (mean, red line) and the 95% confidence interval (shaded).

**Supplementary Fig. 4.**
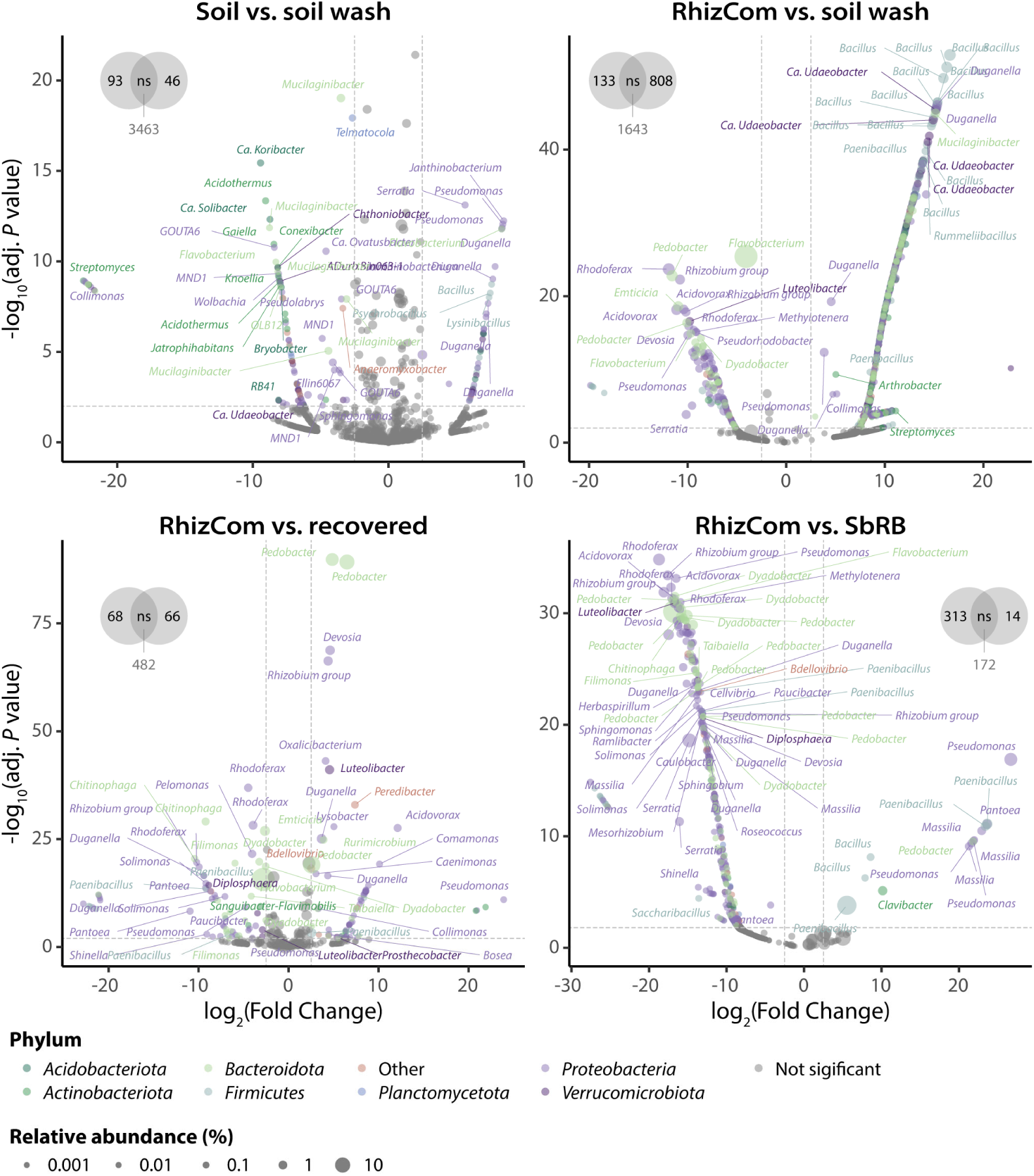
Differential ASV abundance. Differential abundance of ASVs calculated with DESeq2 and using the Wald test with a local estimate of dispersion. ASVs with a |log2(fold change) ≥ 2.5 and an adjusted (adj.) *P* value < 0.01 were considered as significant. Dots represent ASVs, colored according to their phylum and sized according to their relative abundance. ASVs with a relative abundance ≥ 0.05 were named at the genus level, up to a maximum of 45 label overlaps.

**Supplementary Fig. 5.**
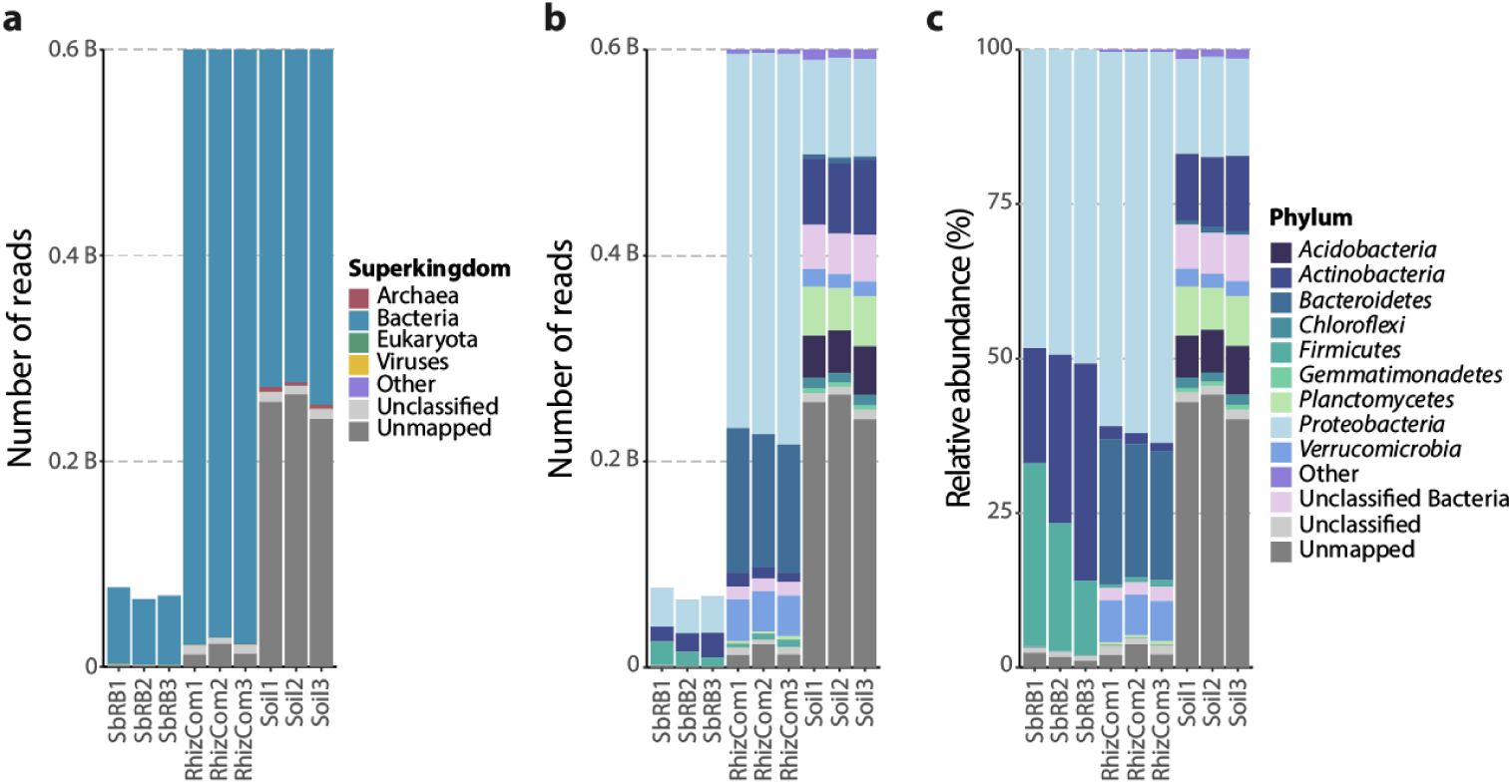
Taxonomic classification of reads in the metagenome samples. **a**, Total number of reads mapping to coding DNA sequences and their taxonomic assignment at the Superkingdom level. **b**, Total number or **c**, relative abundance of reads assigned to Bacteria colored according to their phylum.

**Supplementary Fig. 6.**
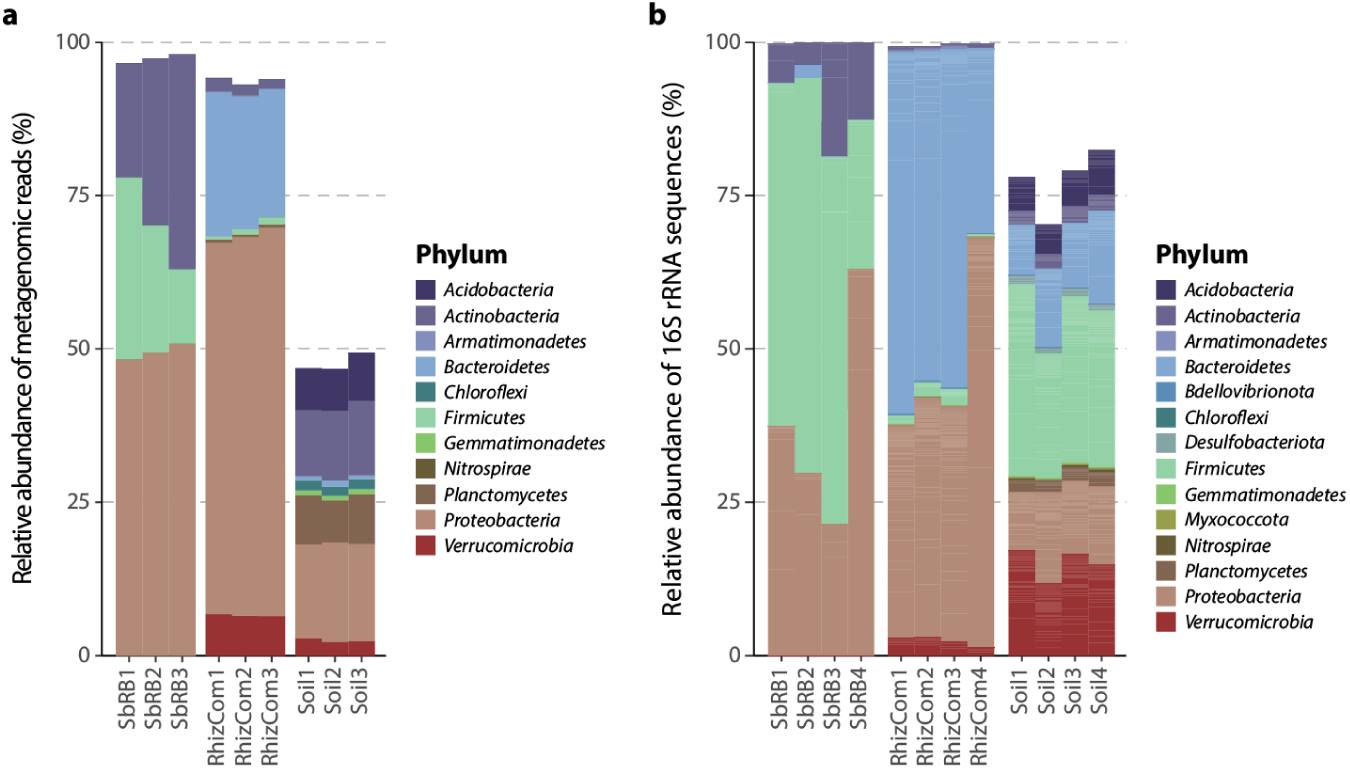
Concordance between taxonomic classification of metagenomic samples vs. samples characterized by 16S rRNA amplicon sequencing. **a**, Relative abundance of metagenomic reads at the phylum level. **b**, Relative abundance of 16S rRNA sequences at the phylum level. Only the top 1,000 ASVs are shown.

**Supplementary Fig. 7.**
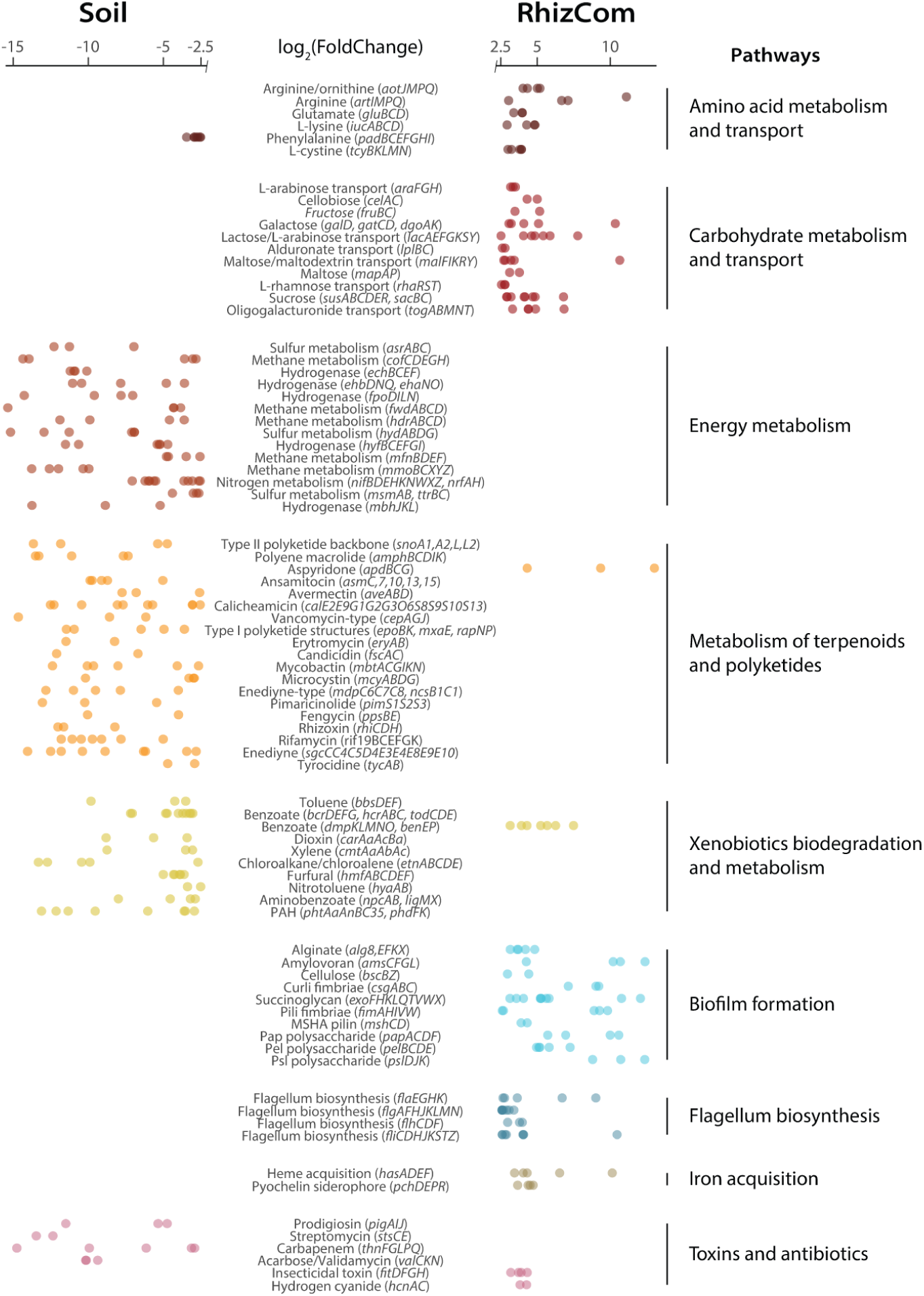
Differential abundance of gene clusters between the RhizCom and soil. Differential abundance of KEGGs annotations between the soil and the RhizCom. Results were filtered to show only significantly enriched annotations (adjusted *P* value ≤ 0.001 and |log2(fold change)| ≥ 2.5) and those annotations that were part of a gene cluster in which multiple genes were differentially abundant. Putative function and gene names are indicated.

**Supplementary Fig. 8.**
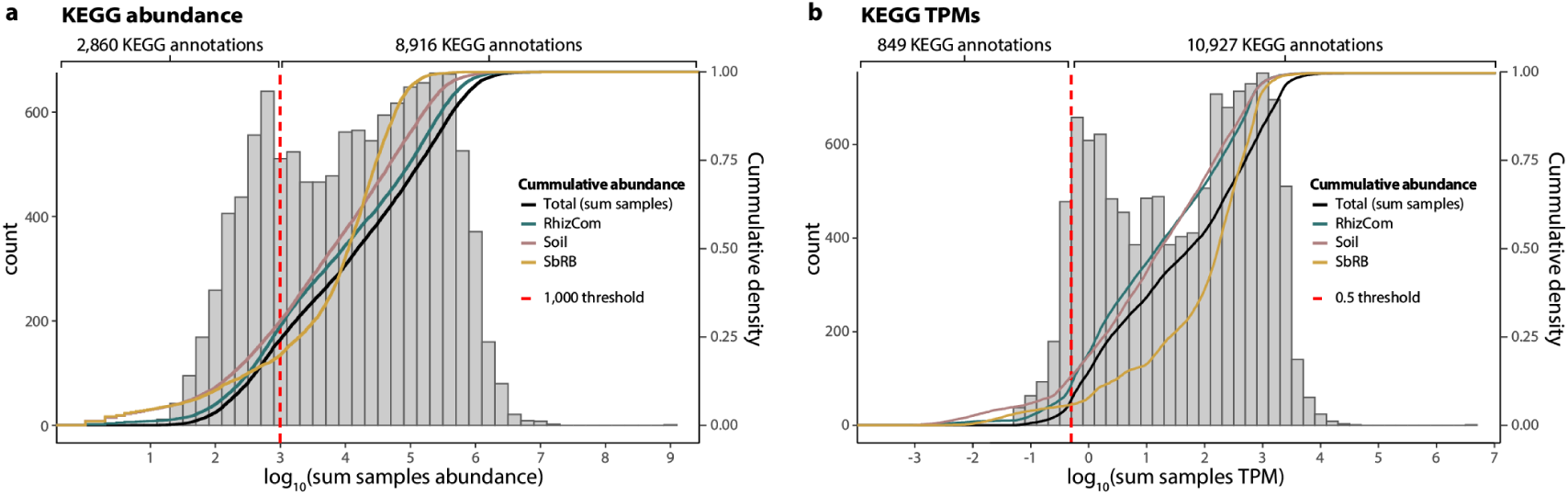
Thresholds of abundance and TPMs for KEGG annotations. **ab**, Histograms (bars, left axis) and empirical cumulative distributions (lines, right axis) of KEGG annotations based on abundance (**a**) and TPMs (**b**). The black line represents the sum of abundances for all samples, while colored lines represent the sum of abundances for the replicates of a sample. The dashed red line indicates an abundance threshold value ≥ 1,000 for KEGG abundances (**a**) or ≥ 0.5 for KEGG TPMs (**b**) used in this study. Numbers above the plot indicate the KEGG annotations above or below the chosen threshold.

## Notes

### Competing Interest Statement

The authors have declared no competing interest.

